# Activation of transposable elements is linked to a region- and cell-type-specific interferon response in Parkinson’s disease

**DOI:** 10.1101/2025.09.03.673956

**Authors:** Raquel Garza, Anita Adami, Arun Thiruvalluvan, Sasvi Wijesinghe, Annabel Curle, Oliver Tam, Talitha Forcier, Danai A Lagka, Nina L Kazakou, Diahann A.M Atacho, Yogita Sharma, Vivien Horvath, Sara Bermudez, Jenny Johansson, Daniel B. Rainbow, Laura Castilla-Vallmanya, Joanne L. Jones, Annelies Quaegebeur, Molly Gale Hammell, Agnete Kirkeby, Roger A. Barker, Johan Jakobsson

## Abstract

Parkinson’s disease (PD) is a neurodegenerative disorder involving a neuroinflammatory response, the cause of which remains unclear. Transposable elements (TE) have been linked to inflammation, but their potential role in PD remains unexplored. Using bulk- and single-nuclei RNAseq of postmortem brain tissue from four brain regions, we studied TE transcription and its correlation with PD neuroinflammation. Over a thousand TEs, including LINE-1s and ERVs, were expressed in a cell-type and region-specific manner in the human brain. Increased TE expression was found in microglia and neurons in the substantia nigra and putamen of PD brains, but not amygdala or prefrontal cortex, compared to controls. This TE activation correlated with an innate immune response in the same brain regions. The link between an interferon response and TE activation was mechanistically confirmed using human pluripotent stem cell-derived microglia and neurons. Our findings provide insights into TE transcription in the PD brain and suggest TEs may contribute to neuroinflammation and pathological progression in PD.

**Teaser:** Transposable elements are linked to a cell type specific inflammatory state in Parkinson’s Disease.

## Introduction

Parkinson’s disease (PD) is an age-related, largely sporadic(*1*) neurodegenerative disorder characterised by, but not limited to, progressive loss of dopaminergic (DA) neurons in the substantia nigra (SN) and the development of alpha-synucleinopathy (a-syn) with Lewy body and Lewy neurite formation(*1*). Emerging evidence from post-mortem, epidemiological, and genetic association studies, along with work in animal models, increasingly supports an important contributory role for neuroinflammation in PD(*2–4*). Notably, features of neuroinflammation, such as microglia activation, occur before neuronal cell death and represent a core component of the pathology early in the disease progression(*2*). However, the mechanism that triggers this inflammatory response remains poorly understood, as do the mechanisms that contribute to its chronicity. A better understanding of how inflammation interacts with and drives disease processes is critical not only to define its role in PD, but also as a potential therapeutic target for this disease.

Transposable elements are viral-like mobile genetic elements that comprise at least 50% of the human genome, including different classes such as Long Interspersed Nuclear Element 1 (LINE-1), Endogenous Retroviruses (ERV), SINE-VNTR-Alu (SVA) and Alus(*5*). Several recent studies have shown that increased TE expression correlates with senescence and other age-related cellular phenotypes, suggesting that aberrant transcription of TEs may be associated with age-related disease processes, such as those seen in neurodegenerative disorders(*6–8*). Age- and disease-related aberrant TE expression is likely to have consequences. For example, it is well known that TEs can act as both transcriptional activators and repressors, affecting the expression of genes in the vicinity of their integration site(*9–12*). TEs appear to be potent as *cis*-regulatory elements in the human brain, where they can act as alternative promoters and enhancers(*11, 13, 14*). Thus, changes in TE activity related to disease and age may disrupt gene regulatory networks by releasing promoter- and enhancer-like activities of TEs in the brain. In addition, aberrant transcriptional activation of TEs may result in the formation of double-stranded RNAs, reverse transcribed DNA molecules, and TE-derived peptides. This can induce ’viral mimicry’, leading to an innate immune system response, including the activation of interferon pathways and other viral response related genes (*15–17*). This type of mechanism has the potential to both induce and maintain a chronic neuroinflammatory response. In line with this, we and others have recently shown that transcriptional activation of TEs in the adult brain causes a neuroinflammatory response in mouse models(*18*). However, our understanding of the potential role of TEs in PD and other neurodegenerative diseases remains limited due to the technical challenges of studying these large repetitive parts of the genome. TEs are composed of highly repetitive sequences that are technically difficult to analyse using standard molecular technologies(*19, 20*). Because of these technical limitations, it is still unclear in which cell and tissue types TEs are expressed, and whether the transcriptional status of TEs differs between the healthy, aged and diseased brain. As a result, the impact of TEs on human physiology, ageing, and disease including neurodegenerative processes remains poorly understood.

In this study, we generated multi-omics data, including bulk- and single-nuclei RNA-sequencing (snRNA-seq) as well as long-read RNA-sequencing from 139 well characterised post-mortem brain samples from Parkinson’s disease patients and sex- and age-matched controls. The analysis concentrated on four brain regions which are differentially affected by PD pathology, reasoning that sampling across this anatomical spectrum would allow us to measure distinct inflammatory states as a function of disease pathology. The tissue selection included the substantia nigra (SN) and putamen (PUT), which are regions known to be majorly affected in PD with loss of the DA-neuron cell body (SN) and the projecting fibres (PUT). In addition, the amygdala (AMY) typically displays major a-syn pathology without major neuronal cell death in PD and the prefrontal cortex (PFC), which only display minor neuronal loss and a-syn pathology in PD. Using a tailored bioinformatics strategy, we quantified TE expression at a locus-specific level per cell type and per brain region. This approach offers an important advantage over existing snRNA-seq bioinformatic strategies, as the regulation and functional effects of a given locus often depends on its precise insertion site, which may be missed by subfamily level analyses.

Our study demonstrates that more than a thousand individual TE loci are expressed in the human brain in a cell-type and region-specific manner, including both LINE-1s and ERVs. In addition, we found increased expression of many TE loci in PD brains which was cell type and region specific, with evidence for transcriptional activation of TEs in microglia in the SN as well as microglia and medium spiny neurons in the PUT. This increased TE expression correlated with the presence of an innate immune response, characterised by the expression of interferon-related and viral response genes, in these brain regions. To prove a direct link between TE expression and an innate immune response, we modelled this *in vitro* using human pluripotent stem cell (hPSC)-derived microglia and dopamine neurons. Our findings provide a unique insight into the complex transcriptional profile of TEs in the human brain which lays the groundwork for investigating the functional role of TEs in PD pathology. Our results not only highlight a role for TEs in the chronic neuroinflammatory state in a neurodegenerative disorder but also provides a bioinformatic approach for the analysis of TEs at single locus resolution, which will enable studies on TEs in other healthy and diseased brain tissues.

## Results

### Single-nuclei and deep bulk RNA-sequencing of human post-mortem Parkinson’s disease tissue across four brain regions

We sampled fresh-frozen post-mortem brain tissue samples from 43 brain donors from the Cambridge Brain Bank, including 25 diagnosed with PD (Braak stages 3-6), as well as 18 sex- and age-matched neurological controls (Figure 1A, Table S1). We obtained a total of 139 brain tissue samples from four different brain regions (SN, PUT, AMY & PFC) which are known to be differentially affected by PD pathology and thus likely to display distinct neuroinflammatory states (Figure 1A). This study design allows for interindividual correlation of region-specific pathological processes and transcriptional alterations, such as TE activation. The tissue was processed for snRNA-seq analysis using the 10X Chromium platform. In total, we sequenced 271,700 high-quality nuclei with an average of 1955 nuclei per sample (Figure S1A), from which we were able to detect an average of 2484 genes (Figure S1A).

**Figure 1.**
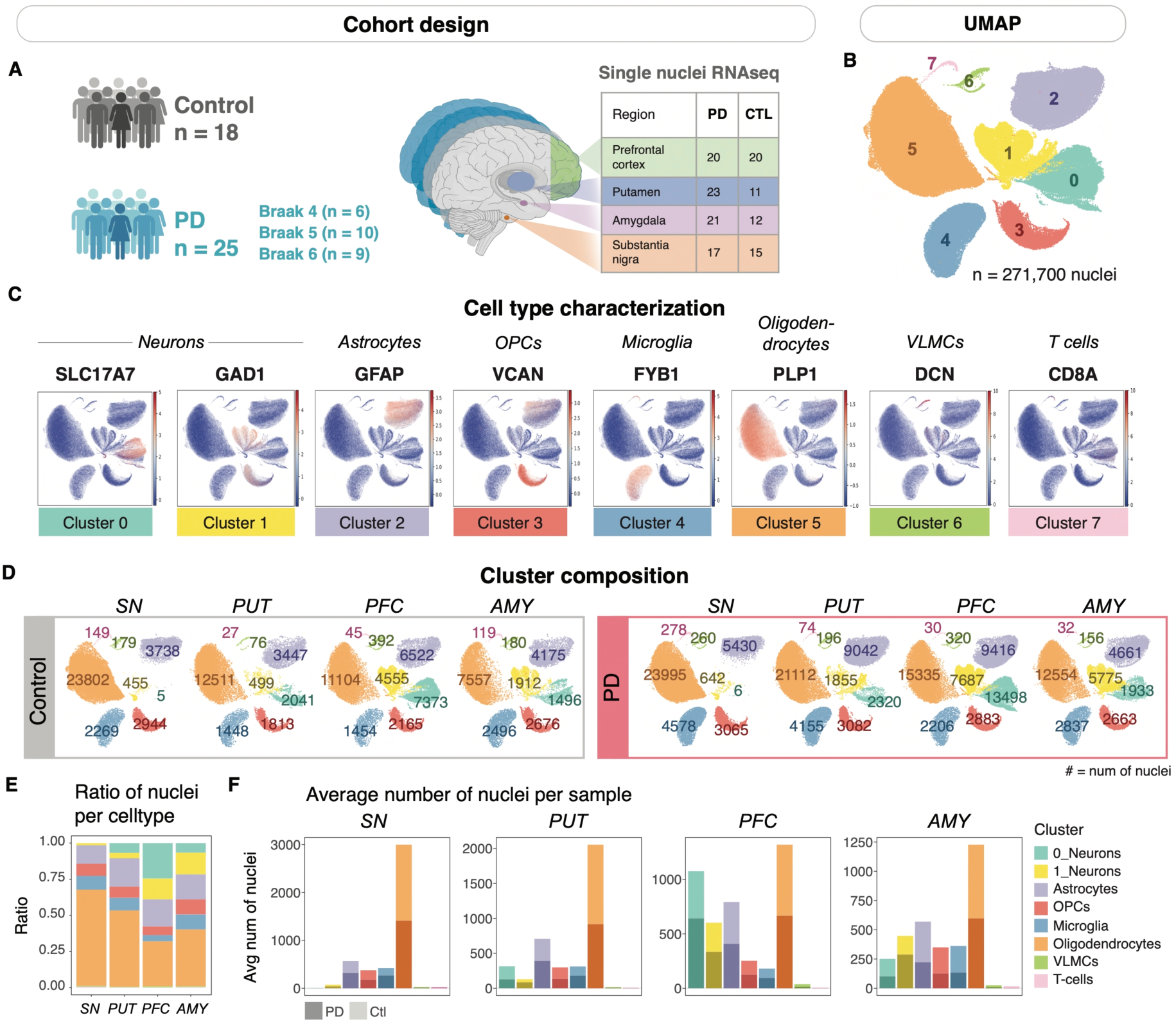
Single-nuclei RNAseq tissue cohort across four regions of PD and control brains. **A)** Schematic showing the cohort design and sample composition. Number of samples sequenced from selected brain regions using snRNA-seq (10X) are listed in the schematic. **B)** UMAP showing unbiased clustering of all sequenced nuclei. The snRNA-seq data cluster into eight defined clusters. **C)** Cell type characterization showing the determined cell type for each cluster and the expression of selected cell type markers. OPC – oligodendrocyte precursor cells, VLMC – vascular leptomeningeal cells. **D)** Cluster composition displaying the number of nuclei in each defined cell cluster and across brain regions and conditions (control vs PD). **E)** Bar plot showing the ratio of defined cell types (y-axis) across the 4 analysed brain regions (x-axis). **F)** Bar plots illustrating the average number of nuclei per sample divided per brain region. Control (ctrl) and PD samples are stacked in the same bar, light colour = control, dark colour = PD.

Unbiased clustering of the snRNA-seq resulted in eight distinct clusters (Figure 1B), which based on the expression of canonical gene markers were classified as neurons, astrocytes, oligodendrocytes, oligodendrocyte precursors (OPCs), microglia, vascular leptomeningeal cells (VLMCs) and T cells (Figure 1C). Neurons generally clustered separately into excitatory neurons (cluster 0), and inhibitory neurons (cluster 1) (Figure 1C, Figure S1B). However, in the PUT, two populations of medium spiny neurons (MSN; DARPP32+/DRD1+/DRD2+) clustered separately, with some MSN present in cluster 0 (150.37 ± 58.54 nuclei per sample (mean ± SE)), and others in cluster 1 (71.33 ± 17.35 nuclei per sample (mean ± SE)) (Figure S1B-C). We found that dopaminergic neurons were present at low numbers in control SN samples and at an even lower number in the PD SN samples (Figure S1D). As expected, the cell type composition varied between brain regions, with PFC displaying the highest ratio of neuronal cells, while the SN tissue was dominated by oligodendrocytes, which is in line with other snRNA-seq studies of these brain regions(*21, 22*). Overall, there was no significant difference in cell type composition when comparing PD and control samples (Figure 1D-F, Figure S1E-G). In summary, this cohort represents a rich resource to study pathological TE expression in the context of neurodegeneration, going beyond the gene-encoding part of the genome.

### Detection of TE expression in the human brain

Most TE loci in the human genome are evolutionarily ancient TE fragments that have lost their regulatory elements and lack the ability to be transcribed. Therefore, we first narrowed down our analysis to evolutionarily young TEs that retain an internal promoter. We focused on full-length LINE-1 elements over 6kb in length and near-complete ERV proviruses, as we and others have found that these elements are expressed in the brain and are implicated in aging and neurodegenerative diseases(*7, 23–30*). To study young TEs at an individual locus level, we performed deep short-read bulk RNA-seq using an in-house 2 × 150 base pair, poly-A enriched stranded library preparation with a reduced fragmentation step to optimise the library insert size for TE analysis on a subset of the post-mortem samples (10 Control, 10 PD, n=65, SN, PUT, AMY, PFC, Figure 2A). Additionally, we sequenced four samples (2 Control, 2 PD) using long-read direct RNA sequencing with Oxford Nanopore technology to further strengthen the observations (919,5 bp average N50) (Figure 2B).

**Figure 2.**
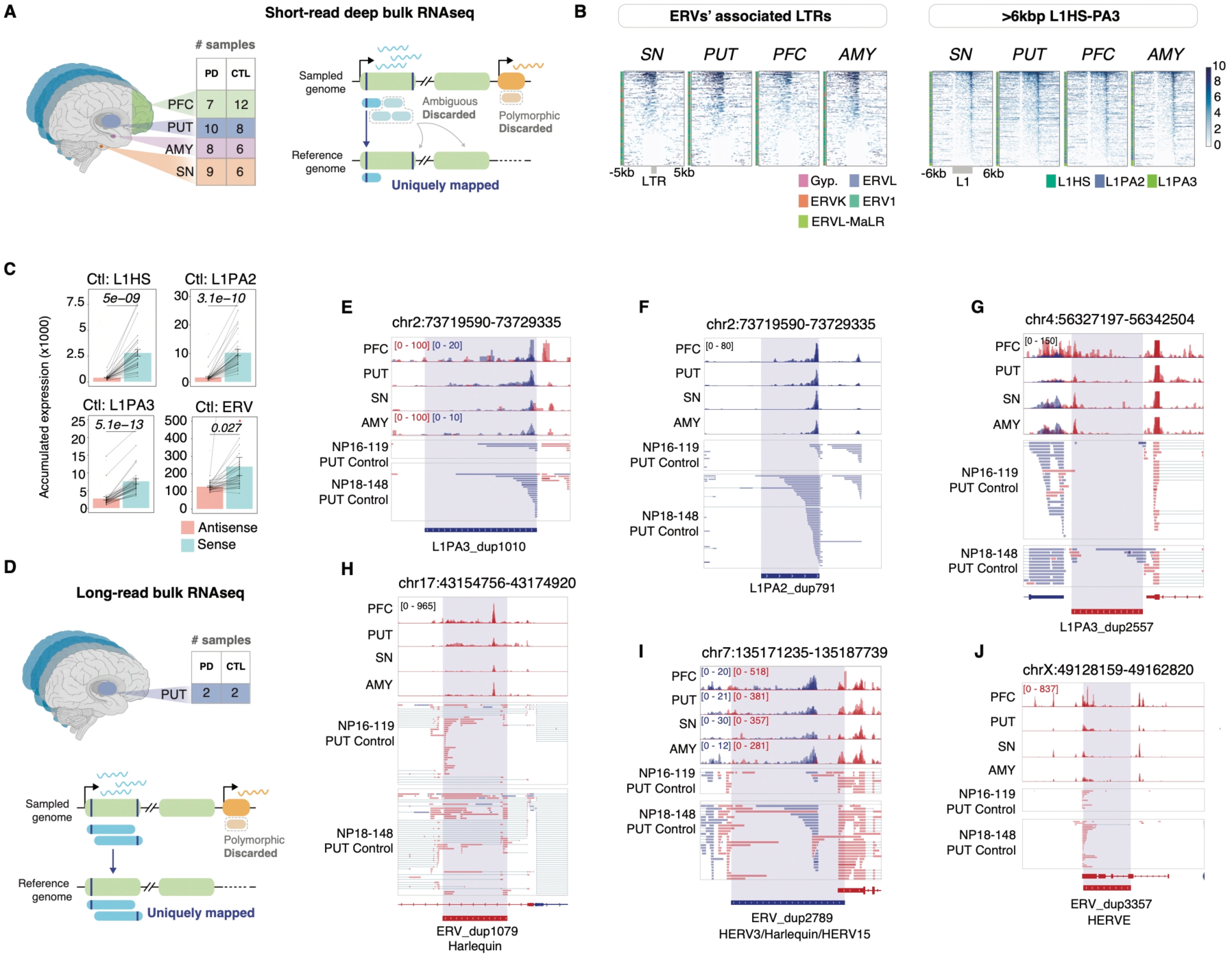
Short and long-read RNAseq detect TE expression through the human brain. **A)** Schematic showing the number of samples analysed using deep short-read RNA-seq and bioinformatical approach to quantify the expression of unique TE loci. **B)** Left: Heatmap showing binned signal (10bp) over the associated LTRs to the expressed ERV elements (from b, n = 739) with 5kbp windows up and downstream. Elements are scaled to 1kbp. Right: heatmap showing binned signal (10bp) over expressed >6kbp L1HS-L1PA3 elements (n = 520) with 6kbp windows up and downstream. Elements are scaled to 6kbp. Color represents log2 RPKM values. **C)** Summed >6kbp young L1 and ERV expression in sense (blue) and antisense (pink) per sample. Bars represent mean normalized expression across samples, error bars show mean ± standard error. **D)** The number of samples sequenced as ONT direct long-read bulk RNA-seq and schematic on long-read mapability. **E-G)** Genome browser tracks showing representative examples of highly expressed >6kbp young L1s. Bulk RNA-seq data on top (range of expression in RPKM scale), ONT direct long-read RNA-seq below. **H-J)** Genome browser tracks showing representative examples of highly expressed ERVs. Bulk RNA-seq data on top, ONT direct long-read RNA-seq below. Blue indicates expression from the positive strand and red expression from the negative strand.

From the deep short read bulk RNA-sequencing, we obtained a total of 32067M reads, with an average of approximately 485M reads per sample. These reads can be mapped uniquely and assigned to individual L1 or ERV loci, except for reads originating from a few of the youngest L1s and polymorphic L1 and ERV alleles not present in the reference genome (Figure 2A). After discarding all ambiguously mapping reads and only quantifying reads mapping uniquely to a single location (unique mapping, Figure 2A) we retained an average of 363M reads per sample (Figure S2A). The TE analysis revealed that many L1 and ERV loci were expressed in both the control and PD brain tissue across all regions. In total we detected 520 L1 and 739 ERV highly expressed loci in the control and/or PD brain, defined as high were elements with more than 10 reads per sample in more than a third of the samples within one of the experimental groups (PD or Control for each region), which were selected for further analyses (Figure 2B). L1 expression was predominantly from primate-specific families, including hominoid-specific elements (L1PA2 and L1PA3) and human-specific elements (L1HS) (Figure 2B). Comparing the number of reads transcribed in the same orientation as the L1s (in sense) to those in the opposite orientation (in antisense) revealed that most of the transcription in these regions was in the same orientation as the L1s (Figure 2C, Figure S2C). The RNA-seq signal over full-length L1s was enriched at the 3′ end, indicating that transcription of L1s terminates at the internal polyadenylation signal (Figure 2B). To verify the transcript structures of these TEs and to further strengthen these observations we sequenced four PUT samples (2 Control, 2 PD) using long-read direct RNA sequencing with Oxford Nanopore technology (Figure 2D). The long-read RNA-seq data confirmed the presence of full-length L1 transcripts that terminate at the internal polyA site (Figure 2E-G). This suggests that most L1 transcripts originate from the L1 promoter and are not a consequence of readthrough or bystander transcription.

To assess the expression of nearly complete ERV proviruses, we used Retrotector predictions, a specialized tool that scans the reference genome to detect LTRs adjacent to ERV internal regions. This makes it possible to distinguish ERV proviruses from solo LTR fragments, which are present in large numbers in the human genome due to recombination events between two endogenous provirus LTRs (*31*). This analysis revealed the expression of many individual ERVs from different lineages in the human brain, including ERV-H, ERV-K, and ERV-W subfamilies (Figure 2B). Due to common recombination events and natural drift since their last retrotransposition proliferation, no ERV in the human genome is intact and many have lost their promoter capabilities. Yet, using the long-read RNA-seq data we could observe transcription initiation from many LTR elements within these ERVs (Figure 2B, H-J). Similar to L1s, transcription of these elements was enriched in the sense orientation (Figure 2C). Notably, ERV expression differs from L1 expression in that several elements display evidence of splicing, as expected from retroviral elements (see e.g., Figure 2J). Additionally, there is readthrough expression to the nearby genome, likely due to a weak poly-A signal in many ERVs (see e.g., Figure 2H).

### Detection of unique TE loci expression at single-cell type resolution in the human brain

The repetitive nature of TEs, combined with the low abundance and stability of TE-derived transcripts, makes quantification of TE expression using single-cell technologies challenging, since this type of data has limited read length and sparse sequencing depth per cell. As a result, snRNA-seq TE analysis is sensitive to noise and signal dropout. We therefore developed a bioinformatic pipeline that leverages the deep bulk RNA-seq to select a subset of TE loci. These elements were then quantified in the snRNA-seq data using a pseudobulk approach per cell type (Figure 3A). For the snRNA-seq analysis we included the 1259 TE loci (n=520 L1s; and n=739 ERVs) which were robustly detected in the bulk-RNA-seq analyses (see Figure 2B).

**Figure 3.**
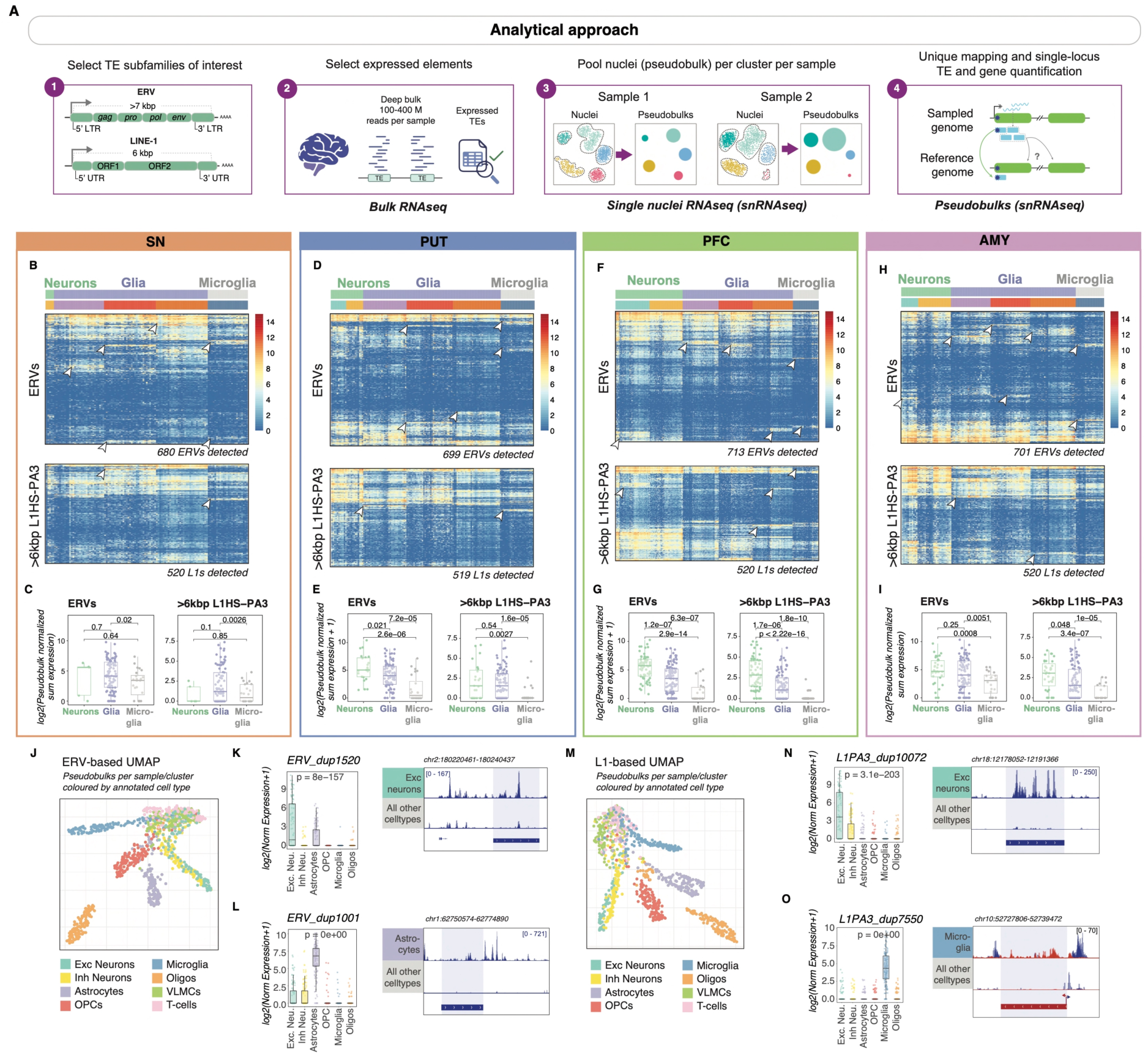
TEs are expressed in a region and cell-type specific manner. **A)** Schematic of analytical approach for cell-type TE quantification. **(B, D, F, H)** Heatmaps showing sample cell-type (columns) TE expression (rows) in the different brain regions. Column annotation showing cell-types. **(C, E, G, I)** Boxplots showing average expression of selected ERVs (top) and >6kbp L1HS-PA3 (bottom) in neurons, glia, and microglia (t-test). Arrows highlighting examples of cell-type specific TE expression. **J)** UMAP dimensionality reduction of pseudobulks (both control and PD) based on ERV expression. Color based on gene-defined cell-types. **(K, L)** Selected examples of cell-type specific ERVs. To the left, boxplots showing cell-type expression of selected ERVs (DESeq2, Wald-test). To the right, genome browser tracks showing cell-type expression (samples overlayed. Normalized by gene size factors). **M)** UMAP dimensionality reduction of pseudobulks (both control and PD) based on >6kbp L1HS-L1PA3 expression. Colour based on gene-defined cell-types. **(N, O)** Selected examples of cell-type specific L1s. To the left, boxplots showing cell-type expression of selected L1s (DESeq2, Wald-test; normalized by gene size factors). To the right, genome browser tracks showing cell-type expression (samples overlayed. Normalized by gene size factors). Genome browser tracks in blue showing transcription from the positive strand, red showing transcription from the negative strand. The centres of all boxplots correspond to the median, hinges correspond to the first and third quartile, and whiskers stretch from the first and third quartile to ±1.5IQR.

To analyze TE expression using snRNA-seq data, our pipeline uses cell-type clusters, previously determined based on gene expression, from each donor. By backtracking the reads from the cells that make up each cluster, we then analyzed the expression of different TE loci using unique mapping in different cell populations from each donor. This pseudo-bulk approach greatly increased the sensitivity of the TE analysis and enabled the quantitative estimation of unique TE locus expression at the resolution of individual cell types across different individuals (Figure S3A-B). Using this approach, we robustly detected the expression of the 1259 TE loci identified in the bulk RNA-seq analysis within the snRNA-seq data (Figure 3B-I, Figure S3C). Conversely, we also confirmed the absence of signal in the snRNA-seq data from ERV and L1 loci that were not expressed in the bulk RNA-seq analysis (Figure S3D). These results confirm that we could reliably detect the expression of a set of transcriptionally active L1s and ERVs in our snRNA-seq data at single cell type resolution.

The TE snRNA-seq pseudobulk analysis revealed a cell-type expression pattern of L1 and ERV loci in both the healthy and PD brain (Figure 3B-I). We found evidence for TE loci that were expressed in all cell types in all brain regions, as well as TEs that were expressed in specific cell-types and specific brain regions (Figure 3B, D, F, H, Figure S3E-F). Across all brain regions we found that both ERVs and L1s were mostly expressed in neuronal cells, followed by ectodermal glial cells, including astrocytes, oligodendrocytes and OPCs, with the lowest expression being seen in microglia. Nevertheless, all cell types displayed L1 and ERV expression and we could detect locus-specific expression in all cell types, including microglia. Notably, UMAP dimensionality reduction demonstrated that it was possible to discern the main cell types in the data based solely on L1 or ERV expression (Figure 3J, M), and differential TE expression analysis of each cell type against all others revealed robust cell-type-specific expression of individual TE loci (Figure 3K-L, N-O), with some showing regional specificity (Figure S3E-F). In summary, these analyses demonstrate that unique L1 and ERV loci are expressed in a cell-type and region-specific manner in the human brain, regardless of disease, suggesting that TE expression is an integral part of the human brain transcriptome.

### Increased TE expression in PD brains

Next, we compared the expression of individual TE loci between PD and control tissue across brain regions and cell types (Figure 4A). We detected a robust increase in expression of many ERV loci in PD microglia in the substantia nigra (SN) and putamen (PUT), but not in the prefrontal cortex (PFC) or amygdala (AMY). We also observed increased L1 expression in microglia in the same brain regions (Figure 4B-D). Other cell types showed substantially less (max 4 upregulated elements) (Figure S4E-F) or no difference in TE expression. We validated these findings using two additional alternative statistical models (see methods “*<u>Validation of</u> <u>differential expression analyses</u>*”; Figure S4G-I). The activated ERVs belonged to several subfamilies, including ERV-K, ERV-W, and ERV-H (Figure S4A). These subfamilies have previously been implicated in neurological disorders(*15, 25, 26, 28, 29, 32*). Many of the activated ERV loci are evolutionary young elements present only in primates and have an almost complete proviral structure, including both LTRs and several putative viral proteins (Figure S4B-D). Some of the upregulated ERVs contain at least one open reading frame of ERV genes (gag, pro, pol, and env). Additionally, we found evidence of ERV activation in a subset of the projecting GABAergic inhibitory MSNs within the putamen (cluster 1) (Figure 4E-F), a cell type which receives a major input from the projecting nigral dopamine neurons. Similarly, we found that these elements represent several families and contain near-complete proviral sequences with the potential to express open reading frames. Our dataset did not contain a sufficient number of dopamine neurons in the SN of PD brains to study this phenomenon in SN neurons, which is not surprising given that these cells represent a small population and are lost early and almost completely in PD.

**Figure 4.**
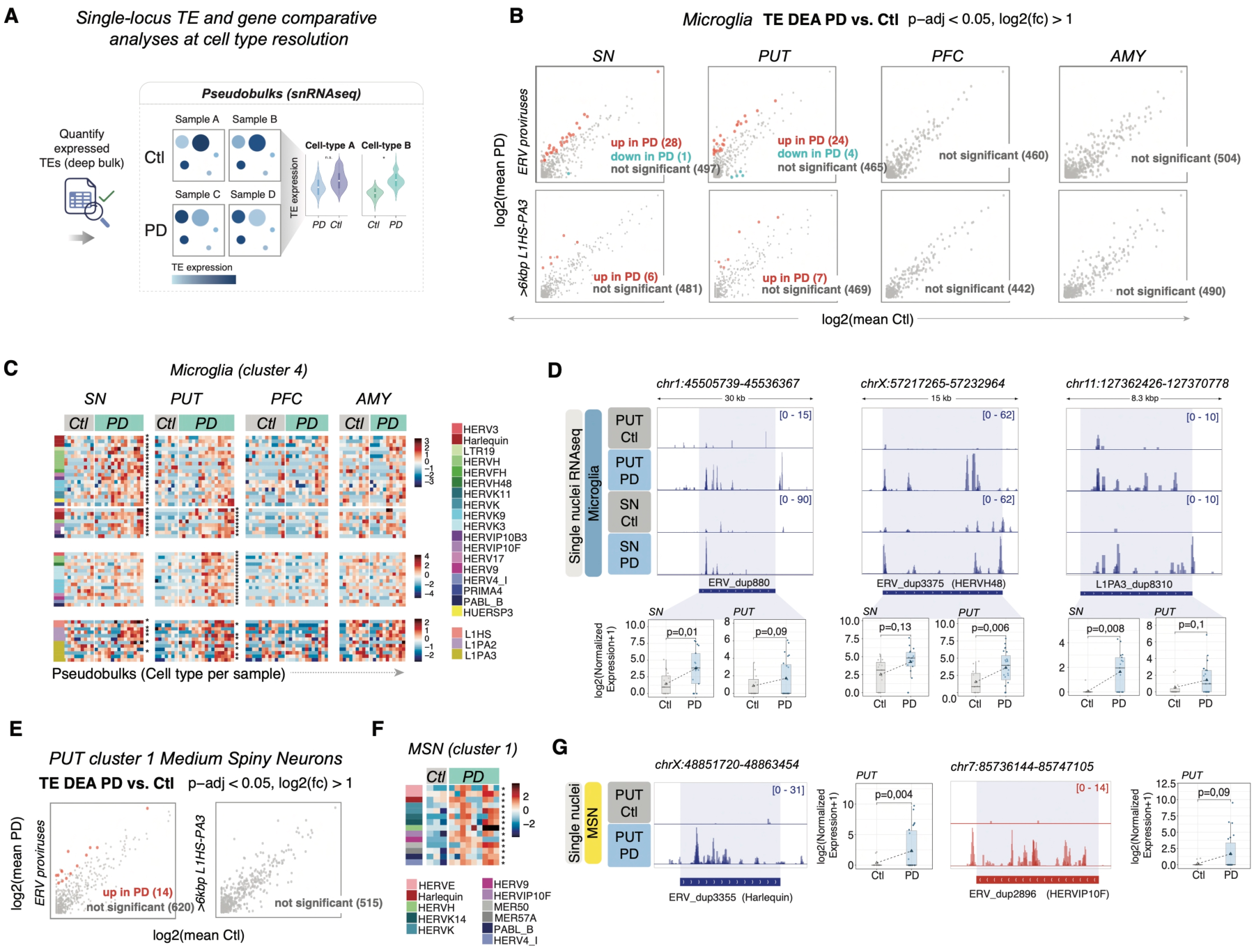
TEs are transcriptionally activated in the PD brain in a region and cell-type specific manner. **A)** Schematic of analytical approach for cell-type TE comparative analysis between PD and control using pseudobulk analysis of the snRNA-seq data. **B)** Mean plots showing ERV (top) and >6kbp L1HS-PA3 (bottom) differential expression analysis results in microglia for each brain region (red indicating significantly upregulated TEs. DESeq2 Wald-test). Num microglia (mean ± SE): Ctl SN = 151.26 ± 32.15, PD SN = 269.29 ± 44.73, Ctl PUT = 131.63 ± 36.65, PD PUT = 180.65 ± 28.88, Ctl PFC = 85.52 ± 16.89, PD PFC = 95.91 ± 17.58, Ctl AMY = 226.9 ± 58.01, PD AMY = 135.09 ± 37.52. **C)** Heatmap showing microglia (column per sample) quantification of upregulated ERVs and >6kbp L1HS-PA3 in SN and PUT (rows). **D)** Top: genome browser tracks showing upregulated ERVK, ERVH48, and >6kbp L1PA3 in PD SN and PUT microglia (forward transcription shown only; normalized by gene size factors). Bottom: boxplots showing quantification in SN and PUT microglia (p-adj, DESeq2, Wald-test; normalized by gene size factors). **E)** Mean plots showing ERV (top) and >6kbp L1HS-PA3 (bottom) differential expression analysis results in cluster 1 of MSNs in PUT (red indicating significantly upregulated TEs (p-adj < 0.05, log2FC > 1). DESeq2 Wald-test). **F)** Heatmap showing PUT MSN cluster 1 (column per sample) quantification of upregulated ERVs in PUT (rows). **G)** Left showing genome browser tracks of upregulated ERVs in PD PUT MSN cluster 1 (Normalized by gene size factors). Right boxplots showing quantification in PUT MSN cluster 1 (p-adj, DESeq2, Wald-test; normalized by gene size factors). Genome browser tracks in blue showing transcription from the positive strand, red showing transcription from the negative strand.

To analyze in detail the TEs that displayed increased expression in the PD brain, we used the long-read direct RNA sequencing data, as this approach has the potential to reveal the complete transcript structure of TEs. In this dataset, we detected 58 of the 73 upregulated TEs (with at least one mapping read, MAPQ > 0). Of these, 53 were ERVs. Analysis of the transcriptional start sites (TSS) of these ERVs revealed that 20 ERVs have a TSS upstream of the element, while 33 initiate transcription from within the element or its immediate proximity. We found evidence of ERVs that initiate expression within the element and at the LTRs (Figure 5A-B). We also found clear evidence that ERVs are spliced, as well as evidence for transcripts encoding putative ORFs (Figure 5B-C). Of the 33 ERVs with TSSs within the element, several serve as TSSs for genes. For instance, we identified an HERVK element on chromosome 2 that functions as the TSS for STAM2, a protein-coding gene involved in the JAK/STAT pathway and associated with frontotemporal dementia(*33*) (Figure 5D). Additionally, we found an ERV on chromosome 13 that serves as a TSS to MPHOSPH8 (MPP8), an epigenetic regulator that is part of the HUSH complex (Figure 5E) (*34*). We also found evidence of ERVs acting as splice sites or transcriptional end sites, including for example HPGDS (Figure 5F), which catalyzes the production of PGD2, that plays an important role in neuroinflammation(*35*) and sleep(*36*). These observations indicate that ERVs upregulated in PD are wired into transcriptional networks. Consistent with this, we found that genes in close proximity to the upregulated ERVs exhibited significantly higher expression in PD microglia and MSNs than in control cells (Figure 5G). Taken together, these data demonstrate that transposable elements (TEs), particularly ERVs, are transcriptionally activated in microglia in the substantia nigra and putamen in patients with Parkinson’s disease, as well as in a subset of MSN in the putamen. Our results suggest that these elements contribute to local gene regulation and may mediate transcriptional alterations associated with PD.

**Figure 5.**
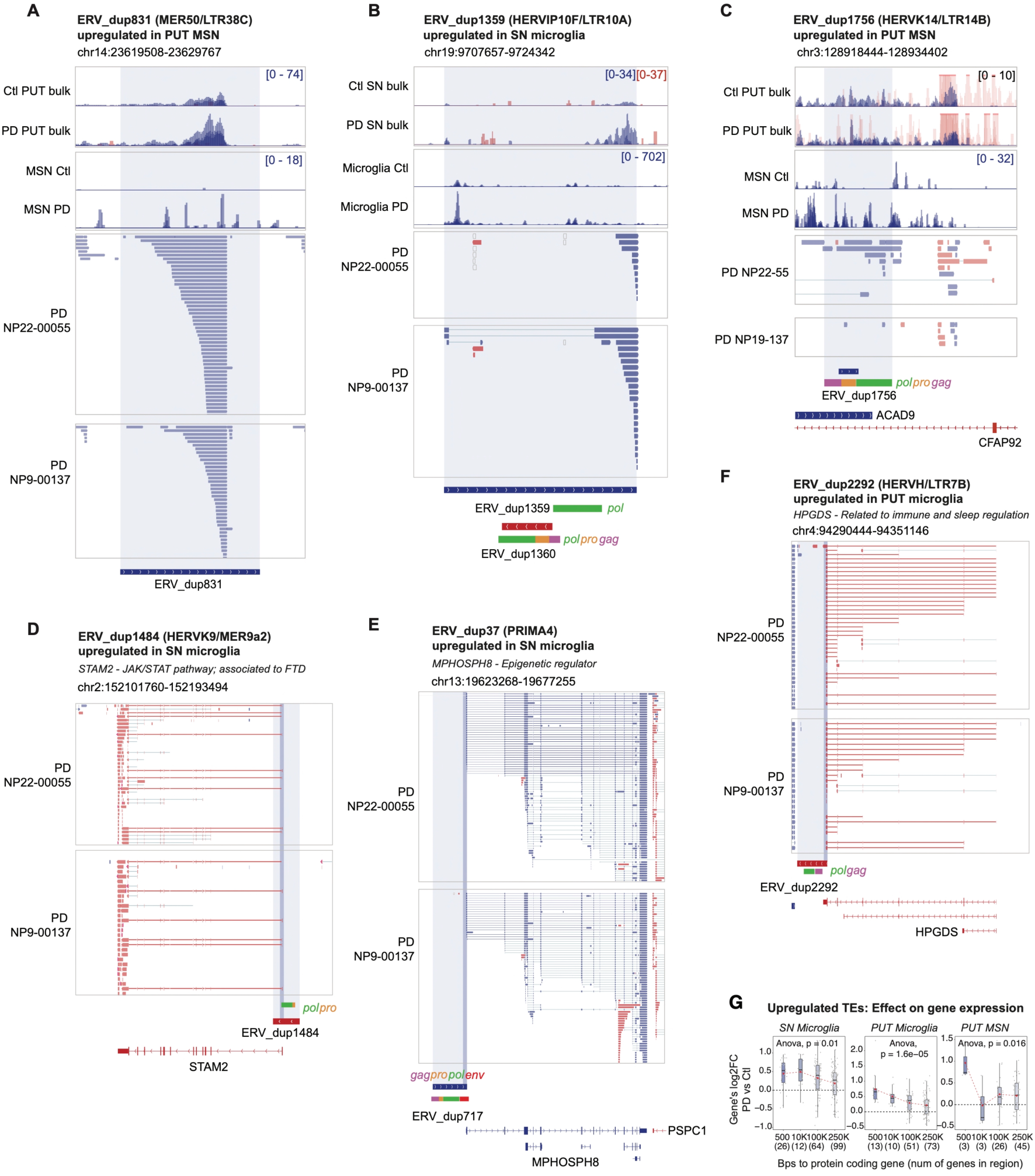
TE expression in the PD brain includes TE-transcripts, chimeric transcripts, and altered nearby gene expression. **A-C)** Genome browser tracks showing examples of upregulated ERVs. ONT direct long-read RNA-seq data of two PUT PD samples. Reads coloured by read strand (blue showing transcription from the positive strand, red showing transcription from the negative strand). **D-F)** Genome browser tracks showing examples of upregulated ERVs hosting transcription start or end sites to protein coding genes. **G)** Boxplots showing change in expression of protein coding genes nearby upregulated TEs (log2FC, PD vs Ctl, Anova). Red dot indicates mean log2FC in window. All boxplot centers correspond to the median, hinges correspond to the first and third quartile and whiskers stretch from the first and third quartile to ±1.5IQR.

### PD microglia display an interferon response that correlates with TE expression

The transcriptional activation of TEs has been linked to innate immunity and interferon responses(*15–17*). Therefore, we investigated whether region-specific TE activation in the PD brain correlates with an interferon response using our snRNA-seq dataset. We found that genes related to the interferon response were upregulated in microglia in the SN and PUT. These genes include toll-like receptors (TLRs) as well as interferon-response factors (IRF), and receptors (IFNGR1) (Figure 6A-B). Notably, the interferon-related genes were not upregulated in microglia residing in PFC or AMY (Figure 6A-B), regions where we saw minimal TE activation in PD (see Figure 4). Gene Set Enrichment Analyses (GSEA) confirmed that, in PD, genes related to terms such as interferon response and viral defense mechanisms were significantly enriched among the upregulated genes in SN and PUT microglia but not in AMY and PFC microglia (Figure 6A-D). These findings were confirmed using two additional alternative statistical models (see methods “*<u>Validation of differential expression analyses</u>*”; Figure S5A-C). The presence of an increased interferon response in the substantia nigra (SN) correlated with increased Braak stage (Figure 6E-F), suggesting that this phenomenon is associated with disease progression or stage in PD. Notably, the activation of an interferon response in PD microglia in SN and PUT coincided with an increase in ERV expression in the same brain regions (Figure 6G). Additionally, we found that the increased expression of certain ERV loci in SN microglia corresponded with an elevated Braak stage, in a similar manner as interferon response genes (Figure 6H).

**Figure 6.**
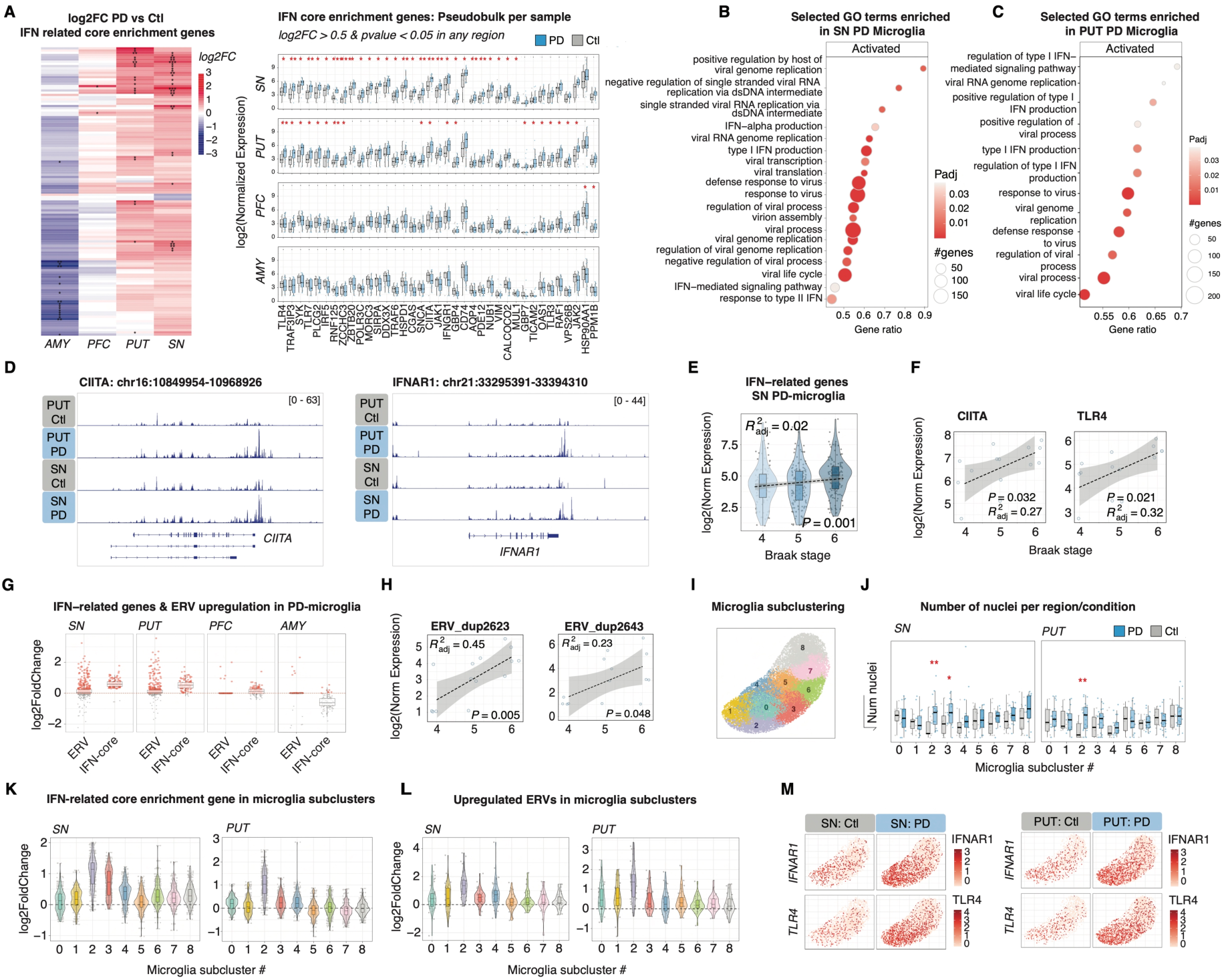
TE activation in PD correlates with an interferon response in microglia. **A)** Left: Heatmap of log2FC (PD vs Control, Wald-test, DESeq2) for genes identified as core enrichment of the GO terms shown in B. Right: Boxplots of samples’ normalized pseudobulk expression in microglia for genes upregulated in PD (log2FC>0.5, p-value<0.05; DESeq2; derived from heatmap). **B-C)** Interferon (IFN) and viral-related GO terms selected from GSEA of PD Microglia (clusterProfiler::gseGO) in SN (B) and PUT (C). **D)** Genome browser tracks of representative genes upregulated in PD SN and PUT microglia. Overlayed tracks show pseudobulk microglial expression from all individuals, normalized by gene sizefactors. Blue=positive-strand, red=negative-strand transcription. **E)** Normalized pseudobulk expression in SN microglia split by Braak stage. Fitted linear model (lm, ggpmisc::stat_poly_line) shown as dotted line with grey standard error. p-value and adjusted R^2^ (ggpmisc::stat_poly_eq). **F)** Linear models (lm, ggpmisc::stat_poly_line) for the association between normalized SN microglial pseudobulk expression and Braak stage for IFN-related genes, with p-value and adjusted R^2^ (ggpmisc::stat_poly_eq). **G)** Boxplots of log2FC (PD vs Control; DESeq2) for all ERV predictions (RetroTector; left) and IFN-related core enrichment genes (as in A). **H)** Linear models (lm, ggpmisc::stat_poly_line) of normalized SN pseudobulk microglial expression and Braak stage for ERVs (examples: ERV17/ERV_dup2623, ERVIP10B3/ERV_dup2643), with p-value and adjusted R^2^ (ggpmisc::stat_poly_eq). **I)** Left: UMAP of microglial subclusters coloured by cluster identity. Right: UMAPs split by region and diagnosis. **J)** Boxplot of nuclei number per individual per cluster (y-axis: squared root of number of nuclei). T-tests show significant differences in SN (clusters 2 and 3) and PUT (cluster 2). **K-L)** Violin plots per microglia subcluster showing log2FC of IFN-related core enrichment genes (K) and upregulated ERVs (L) in SN and PUT. **M)** UMAP showing expression of representative genes upregulated in clusters 2 and 3. Boxplots: center=median; hinges=first and third quartile; whiskers extend to ±1.5IQR.

To further investigate transcriptional changes in microglia, we subclustered the microglia population for increased resolution of subpopulations. This resulted in nine distinct clusters (Figure 6I). We found that clusters 2 and 3 were significantly enriched for cells from PD brains, particularly microglia from the substantia nigra (SN) and putamen (PUT) (Figure 6I-J, Figure S5D). Notably, these two populations were characterized by an increased expression of interferon genes, such as TLRs and IRFs, as well as an increased expression of ERVs (Figure 6K-M). Taken together, these results demonstrate that a disease-specific population of microglia exists in the SN and PUT of PD brains that is characterized by an innate immune response and increased TE expression.

### Interferon treatment results in activation of TEs in human microglia and dopamine neurons

To mechanistically investigate the link between interferon activation and ERV expression, we performed *in vitro* experiments using human microglia. We differentiated human pluripotent stem cells (hPSCs) into microglia using a stepwise protocol involving mesodermal induction via embryoid body formation, followed by hemogenic endothelium induction and myeloid differentiation to generate primitive monocyte-like precursors(*37, 38*). These precursors were then matured into microglia by supplementing with key neuron-derived factors that promote microglial identity, thereby recapitulating human microglia in monoculture (Figure 7A). The differentiated cells displayed morphological features and transcriptional signatures of human microglia (Figure 7B-D).

**Figure 7.**
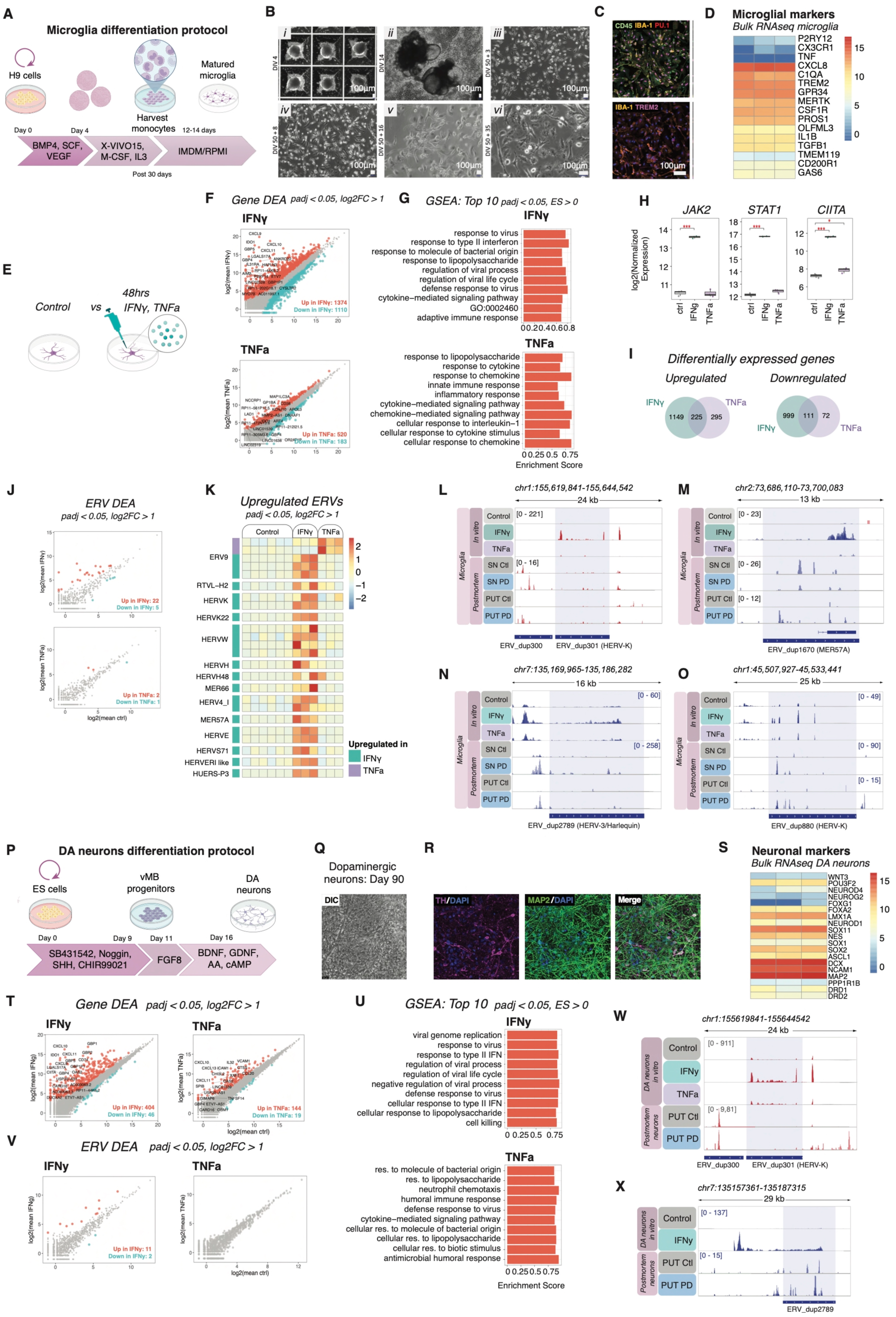
Interferon treatment in human microglia and dopaminergic neurons leads to increased TE expression. **A)** Schematic of human microglia differentiation protocol. **B)** Brightfield images of differentiation stages (DIV=days in-vitro differentiation). Scale bar=100μm. **C)** Immunostaining for microglial markers. Scalebar=100μm. **D)** Heatmap of microglial markers (bulk RNA-seq). **E)** Schematic of cell culture treatments. **F)** Mean-plots of gene differential expression analysis (DEA) in untreated vs IFNγ- and TNFa-treated microglia (DESeq2, Wald-test). Differentially expressed (DE) genes highlighted in red (log2FC>1, padj<0.05) and blue (log2FC<-1, padj<0.05). **G)** Top-10 enriched GO-terms in IFNγ- and TNFa-treated microglia (GSEA, clusterProfiler). GO:0002460=adaptive immune response based on somatic recombination of immune receptors built from immunoglobulin superfamily domains. **H)** Normalized expression of selected genes in untreated (ctrl) and treated microglia (DESeq2, Wald-test). Center=median; hinges=first and third quartile; whiskers extend to ±1.5IQR (*padj <0.05, **<0.01, ***<0.001) **I)** Venn-diagrams of DE-genes across treatments (padj<0.05, upregulated: log2FC>1, downregulated: log2FC<-1). **J)** Mean-plots of ERV DEA in untreated vs IFNγ- and TNFa-treated microglia (DESeq2, Wald-test), colored as F. **K)** Heatmap of DE ERVs across treatments. Annotation indicates experiment of upregulation; labels show overlapping LTR families (Repeatmasker) for each ERV provirus. **L-O)** Genome browser tracks of ERVs upregulated in IFNγ-treated microglia across treatments and in postmortem SN and PUT microglia pseudobulks. **P)** Human dopaminergic (DA) neuron differentiation protocol. **Q)** Brightfield and **R)** Immunostainings of DA neurons. **S)** Heatmap of neuronal markers expression (bulk RNA-seq). **T)** Mean-plots of gene DEA in untreated vs IFNy- and TNFa-treated DA neurons (DESeq2, Wald-test), colored as in F. **U)** Top-10 GO-terms in IFNy- and TNFa-treated DA neurons (GSEA, clusterProfiler). **V, X)** Genome browser tracks of ERVs upregulated in IFNy-treated DA neurons across treatments and in postmortem MSN PUT pseudobulks (clusters 0 and 1). Blue=positive-strand, red=negative-strand transcription. **W)** Mean-plots of ERV DEA in treated DA neurons (DESeq2, Wald-test), colored as in F.

To model the interferon response detected in PD microglia in the substantia nigra (SN) and putamen (PUT), we treated mature hPSC-derived microglia(*37, 38*) with interferon-gamma (IFNγ; 5 ng/mL; 48 hrs) (Figure 7E). To evaluate whether TE upregulation is specific to an IFNγ response, we also treated the microglia with a different immunostimulant, tumour necrosis factor-α (TNFα; 5 ng/mL). TNFα a pleiotropic cytokine that also regulates homeostatic functions in the central nervous system(*39*). Following 48 hours stimulation, microglia were harvested for 2 × 150 bp paired-end, strand-specific bulk RNA-seq analysis.

Transcriptomic analysis revealed stimulus-specific responses (Figure 7F-G). IFNγ-treatment led to a typical type II interferon response, including robust activation of JAK/STAT signaling (JAK2:log2FC 3.06, padj 7,93e-238; STAT1: log2FC 4.69, padj < 2.22e-308) as well as downstream immune-related genes (Figure 7H) including the MHC class II transactivator - CIITA (log2FC 4.38, padj 1,76e-291), which plays a key role in regulating Class II expression (Figure 7H). In line with this, gene set enrichment analysis revealed that IFNγ treatment results in the transcriptional activation of genes linked to antiviral response terms (Figure 7F-G). TNF-α treatment also resulted in the activation of immune-related transcriptional programs; but without JAK2-STAT1 upregulation (Figure 7G-H) or enrichment of antiviral signatures, indicating engagement of alternative innate immune pathways (Figure 7H-I). Thus, the two treatments resulted in distinct inflammatory responses and are well suited to study the specificity of the TE upregulation in relation to an interferon response in microglia, as found in the SN and PUT of PD brains (Figure 6B-C).

Analysis of TE expression in the immune-stimulated microglia revealed that many TEs, in particular loci belonging to several different ERV lineages, were robustly upregulated by IFNγ-treatment (Figure 7J-O). TNFα treatment, however, had a very limited effect on ERV expression (Figure 7J-O), suggesting specificity of ERV expression to interferon related pathways. Notably, the upregulated ERV loci upon IFNγ treatment included several almost intact provirus insertions, which are evolutionary young elements with the potential to be reverse transcribed as well as be translated to produce ERV-derived peptides. We also noted that some of the most robustly upregulated provirus insertions upon IFNγ treatment were also found to be upregulated in PD microglia in the SN and PUT (Figure 7L-O). We validated the increased expression of ERV upon IFNγ treatment using microglia derived from an independent human PSC cell line (Figure S6).

Since we also detected increased TE expression in neurons in the PD brain, we differentiated hPSCs into midbrain dopamine neurons using our previously established protocol (Figure 7P)(*40*). The differentiated cells displayed the morphology of mature human dopamine neurons and expressed marker genes characteristic of this cell type, such as TH and MAP2 (Figure 7Q-S). We treated the DA neurons with IFNγ and TNFα using an approach similar to that used for the microglia treatment experiments. Like the microglia, IFNγ treatment of the dopamine neurons resulted in a robust transcriptional response, including the activation of gene programs related to innate immunity, interferon response, and antiviral defense (Figure 7T-U). TNFα stimulation induced the activation of immune-related gene programs, but it was not associated with antiviral immune responses observed with IFNγ (Figure 7T-U). In DA neurons, we found that IFNγ treatment resulted in the transcriptional activation of TEs (Figure 7V), while TNFα treatment did not (Figure 7V). The IFNγ-activated TEs in DA neurons were ERVs (Figure 7V-X). Notably, most of the individual loci that were upregulated in DA neurons (total of 11) were distinct from the ERVs activated in IFNγ-treated microglia (7 unique upregulated ERVs in DA neurons). This is similar to what we found in the putamen of the PD brain, where the interferon response correlated to the activation of ERVs in both microglia and neurons, albeit different loci (Figure 7V, Figure 4B, E). Taken together, these results demonstrate that activation of an IFN response in human microglia and DA neurons leads to TE transcriptional activation. This provides a mechanistic explanation for our observations of a correlation between an interferon response and TE expression in the PD brain. Notably, while both neurons and microglia activate TE expression, the individual loci that are activated differ, suggesting that the downstream consequences of this transcriptional response are cell-type dependent.

## Discussion

At least 50% of the human genome consists of transposable elements (TEs), which have colonized our germline throughout evolution(*5*). It has been thought that TEs are transcriptionally silenced via epigenetic mechanisms, including DNA and histone methylation, in adult tissues(*41, 42*). However, emerging data from our laboratory and others have begun to challenge this view by demonstrating that some TEs are selectively expressed in various cell types in the mammalian brain, where they could play an important regulatory role in both health and disease(*13, 43–47*). Several previous studies have found elevated TE expression in various neurodegenerative diseases including multiple sclerosis(*29, 48*), amyotrophic lateral sclerosis(*25, 26*), as well as in surgical biopsies after traumatic brain injury(*47*). However, the abundance and repetitive nature of TEs make it difficult to accurately measure their transcription, particularly at the resolution of individual loci(*19, 49*). Thus, most of these recent studies have considered TEs as part of different TE subfamilies, such as L1PAs or groups of ERVs, thereby limiting our understanding as to how many TEs are expressed in the human brain and in which cell types they are expressed. In this study, we addressed this issue by using a combination of deep bulk and snRNA-seq with custom bioinformatics pipelines. Leveraging our deep bulk RNA-seq data, we informed our cell-type specific TE quantification to ignore false positives that could arise due to mispriming or pre-mRNA content in 10X libraries(*50*). Our approach draws on the strength of each technology: deep bulk RNA-seq, providing sensitive identification of TE-transcripts, and snRNA-seq providing cell type specific information. Notably, we were able to capture many TE-transcripts using long-read direct RNA, which independently confirms the expression of these TEs in the human brain. However, our approach also has shortcomings. For example, these findings likely extend to other TE subfamilies, such as Alu and SVAs. However, 10X libraries do not provide a robust readout for these subfamilies, as elements detected in bulk RNA-seq were not captured by the 10X snRNA-seq, likely due to their low mapability or other technical shortcomings related to transcript amplification. In summary, our approach demonstrates that more than a thousand unique L1 and ERV loci are expressed in the adult human brain in a highly cell-type-specific manner. Our data shows that evolutionarily young TEs substantially contributes to the human transcriptome and provide direct evidence, with unprecedented detail, that TEs are transcriptionally activated in PD.

We further found that TE expression was upregulated in microglia in the putamen and substantia nigra as well as in a subset of medium spiny neurons in the putamen. The majority of the upregulated TEs in PD were ERVs, which are remnants of old retrovirus infections that have entered our germline and make up about 8% of the human genome. Previous studies using human cell culture models have demonstrated a strong *cis*-regulatory effects of ERVs related to immune responses. These studies suggest that ERVs play a key regulatory role in immune-related genes by acting as enhancers or start, end, or splice sites(*51–53*). In this regard, our results highlight the importance of locus- and cell-type-specific analyses. Several of the ERVs that we found to be upregulated in PD acted as transcriptional start sites (TSS) or stop sites for protein-coding genes, demonstrating that ERVs are wired into regulatory networks that are altered in PD. Due to cell-type heterogeneity and the relatively sparse presence of microglia in these samples, the isoforms of these TE-fusion genes are likely underestimated in our analysis. Follow-up studies using high-throughput single-cell long-read sequencing technologies, as well as mechanistic studies, are crucial for understanding the relevance of these transcripts in the context of the disease.

It is worth noting that several of the proviral loci activated in microglia or striatal medium spiny neurons in the PD brain contain near-intact proviruses, many of which have the capacity to generate RNA/DNA hybrids as well as peptides. In *Drosophila* or mouse models, there is growing evidence linking the upregulation of ERVs and other TEs to neuroinflammation and neurodegeneration(*7, 18, 54*). Several studies have shown that aberrant ERV expression results in the formation of double-stranded RNAs and reverse-transcribed DNA molecules, which can trigger an innate immune response via viral mimicry(*16, 17, 55*). Emerging data also indicates that ERV transcriptional activation results in the production of peptides that can activate immune pathways(*16*), or lead to the presentation of ERV-derived antigens(*55, 56*). Additionally, aberrant ERV-derived peptide expression has been reported to impair synaptic function and contribute to protein aggregation(*24, 54, 57*), processes highly relevant to PD pathology and other neurodegenerative disorders. Thus, the transcriptional activation of TEs in the PD brain is likely to have several consequences that contribute to PD pathology, including the dysregulation of gene regulatory networks, boosting or sustaining the neuroinflammatory response as well as impacting on synaptic function and protein aggregation. However, the development of robust reagents and protocols to identify ERV-derived molecules with specificity to unique loci will be key to clarifying how disease-associated ERVs are involved in neuroinflammatory responses and neurodegenerative states.

Evidence from neuropathological and neuroimaging studies supports the presence of an early, chronic neuroinflammatory process in PD(*58*) but the triggers and drivers of this are unclear. In this study, we found that molecular changes in microglia differ in different brain regions in PD, even when the tissue samples come from the same individual. Within the nigrostriatal pathway, we identified a population of microglia characterized by activation of the interferon response and antiviral gene programs, but we did not see these responses in the amygdala nor prefrontal cortex despite both sites showing varying degrees of alpha synuclein pathology. Thus, our results demonstrate that the transcriptional activation of TEs correlates with an interferon response in the same brain regions, but not necessarily with the degree of a-synuclein pathology. Our mechanistic *in vitro* modeling, which uses hPSC-derived microglia and DA neurons, confirmed that inducing an interferon response causes transcriptional activation of TEs. Therefore, TE transcriptional activation appears to be downstream of the interferon response. Currently, we do not understand why an IFN response results in the activation of ERVs, but it has been suggested that IFN-relevant transcription factors such as STAT proteins directly bind to TEs, including ERVs(*53*). Investigating the transcription factors recruited to TEs upon IFN treatment and whether this results in chromatin remodeling that releases their transcriptional silencing will be interesting. It is worth noting that the TE loci activated in microglia and neurons, both *in vivo* and *in vitro*, were distinct for each cell type, indicating that underlying epigenetic states play an important role in TE activation. Additionally, we note that the amplitude of the IFN response as well as the expression of some ERVs increases with increasing Lewy Body pathology as defined by Braak stages, which supports a link to the ongoing disease process, although exactly how is unknown. Indeed, the initial trigger of the IFN response in PD remains unclear. Our *in vitro* modeling experiments cannot determine whether TE expression activation or the IFN response is the cause or consequence of the other in actual PD brains. The trigger of the IFN response may involve TEs, but may also be linked to the release of mitochondrial DNA into the cytosol or genomic DNA damage. These processes all increase with aging, which is the main risk factor for PD. More studies mechanistically dissecting the role of TEs in inflammation will determine whether TEs can act as a trigger or play into a feed-forward loop stimulating the activation of inflammatory pathways.

In summary, we here describe the presence of an IFN response in microglia the substantia nigra and putamen that is mechanistically linked to the transcriptional activation of TEs in the PD brain, thus suggesting a role for TEs in the pathogenesis of PD. A finding that could lead to new treatment avenues for PD and other related chronic neurodegenerative disorders.

## Methods

### Brain tissue sampling

Human post-mortem brain tissue from donors with Lewy Body pathology and neurologically healthy controls was sourced from the Cambridge Brain Bank with approval from the London– Bloomsbury Research Ethics Committee (REC reference no. 16/LO/0508). The Cambridge Brain Bank is fully licensed by the Human Tissue Authority, a government regulatory body which oversees consent, storage, and the use of human samples in research (under HTA PM License 12318). Donors with Lewy Body pathology (n=25) had a clinical diagnosis of Parkinson’s disease and were compared to an age- and sex-matched group of neurologically healthy control donors (n=18) who had no history of neurological illness. A neuropathologist assessed all cases for Lewy Body pathology Braak staging, co-existing proteinopathies and other pathologies. Fresh frozen tissue was sampled from SN, PUT, AMY, and PFC at the level of Brodmann area 46 (PFC).

### Nuclei isolation from post-mortem brain tissue

The nuclei isolation from sampled post-mortem brain tissue was performed as previously described(*18, 59*) using a sucrose gradient-based isolation. Briefly, a small section of the tissue (3 mm^3^) was cut from the sample tissue and dissociated in 1 ml of ice-cold lysis buffer (0.32 M sucrose, 5 mM CaCl_2_, 3 mM MgAc, 0.1 mM Na_2_EDTA, 10 mM Tris-HCl, pH 8.0, 1 mM DTT) using a 1 ml tissue douncer (Wheaton). The lysate was then carefully layered on top of a 2.8 ml sucrose cushion (1.8 M sucrose, 3 mM MgAc, 10 mM Tris-HCl pH 8.0, and 1 mM DTT diluted in milliQ water) and then centrifuged at 30,000 × g for 2 hours and 15 min. Once the supernatant was removed, the pelleted nuclei were softened for 10 min in 50 μl of nuclear storage buffer (15% sucrose, 10 mM Tris-HCl pH 7.2, 70 mM KCl, and 2 mM MgCl_2_, all diluted in milliQ water), resuspended in 300 μl of dilution buffer (10 mM Tris-HCl pH 7.2, 70 mM KCl, and 2 mM MgCl_2_, diluted in milliQ water) and then filtered through a cell strainer (70 μm). The nuclei were sorted via fluorescence-activated nuclear sorting (FANS) with a FACS Aria (BD Biosciences) at 4° C at low flow rate using a 100 μm nozzle (reanalysis showing >95% purity).

A detailed protocol can be found at doi: dx.doi.org/10.17504/protocols.io.5jyl8j678g2w/v1.

### Single-nuclei RNA sequencing

The nuclei for single nuclei RNA sequencing (8500 nuclei per sample) were loaded onto the Chromium Next GEM Chip G Single Cell Kit along with the reverse transcription mastermix according to the manufacturer’s protocol for the Chromium Next GEM single cell 3’ kit (10X Genomics, PN-1000268) to generate single-cell gel beads in emulsion. cDNA amplification was achieved following the guidelines from 10X Genomics employing 13 cycles of amplification of the 3’ libraries. Sequencing libraries were generated with unique dual indices (TT set A) and pooled for sequencing on a Novaseq6000 or Novaseq X plus using a 100-cycle kit and 28-10-10-90 reads. A detailed protocol can be found at dx.doi.org/10.17504/protocols.io.kxygx41ndl8j/v1.

### Bioinformatic analysis of single-nuclei RNA sequencing

Basecalling and sample-specific fastq files were generated using 10x Genomics mkfastq(*60*) (version 6.0.0; RRID:SCR_017344). Mapping and gene quantification per nuclei was performed using 10x Genomics cellranger count (version 6.0.0) including introns (--include-introns) and using hg38 (GRCh38) as the reference genome.

#### Quality control

Samples identified as hypoxic during pathological examination were excluded from further analysis (individuals NP17-94 (4 samples) and NP22-00075 (4 samples)). Selection of nuclei and features for downstream analysis was performed using Scanpy (version 1.10.1; RFID). Read count matrices from cellranger were loaded to sample-specific anndata objects using Scanpy (scanpy.pp.read_10x_mtx).

Genes expressed in less than three nuclei were excluded from further analysis (scanpy.pp.filter_genes, min_cells = 3). Quality control metrics were calculated using Scanpy’s calculate metrics function using mitochondrial and ribosomal genes (scanpy.pp.calculate_qc_metrics, percent_top = None, log1p = False). Nuclei were discarded from further analyses if any of the following conditions were met:

1. It was identified as a doublet using Scanpy’s scrublet function on default parameters (scanpy.pp.scrubblet).
2. Had more than 2% of mitochondrial counts.
3. Had less than 500 genes detected.
4. Had a higher number of genes detected than the mean plus two standard deviations in the sample, or under the mean minus two standard deviations.

Furthermore, any sample retaining fewer than 50 nuclei following the aforementioned filtering steps was excluded from further analysis. In total, the downstream analysis was performed in 139 samples.

#### Clustering and UMAP projection

Samples were merged using anndata’s concatenation function (adata.concat, join = outer). Normalization of counts was performed using Scanpy’s normalization function (scanpy.pp.normalize_total, target_sum = 1e6), natural-log transformed with a pseudocount of 1 (sc.pp.log1p) and scaled to a max value of 10 (scanpy.pp.scale, max_value = 10).

Principal component analysis was performed with an arpack solver to account for the sparsity of the matrix (scanpy.tl.pca, svd_solver = arpack). Nearest-neighbor distances calculations, and neighborhood graph were made using 20 principal components, and a neighborhood size of 15 (scanpy.pp.neighbors, n_neighbors = 15, n_pcs = 20). Clustering was performed using the leiden algorithm (igraph implementation) with a resolution of 0.1 (scanpy.tl.leiden, directed = False, flavor = igraph).

A graph abstraction was built using Scanpy’s partition-based graph abstraction (PAGA) implementation (scanpy.tl.paga) prior to run the Uniform Manifold Approximation and Projection (UMAP) (scanpy.tl.umap, init_pos = paga).

Harmony was used to integrate samples across sequencing batch, sex, and individuals (scanpy.external.pp.harmony_integrate). Nearest-neighbor distances were recalculated using the resulting PCA (scanpy.pp.neighbors, n_neighbors = 15, n_pcs = 20, use_rep = X_pca_harmony). Clustering was then performed using the leiden algorithm (scanpy.tl.leiden, resolution = 0.1, directed = False, flavor = igraph). A paga graph abstraction was built (scanpy.tl.paga) and used to run the UMAP (scanpy.tl.umap, init_pos = paga).

#### Cell type characterization

Cell cycle scores were calculated with a previously reported list of cell-cycle related genes(*61*) using Scanpy’s cell cycle scoring function (scanpy.tl.score_genes_cell_cycle).

Marker genes were obtained per region, per cluster, using Wilcoxon tests (scanpy.tl.rank_genes_groups, group_by = leiden, method = wilcoxon). Cell type annotation was manually done upon examination of the resulting list of marker genes and canonical cell type markers.

#### Gene and TE expression quantification using pseudobulks

A file per sample specifying barcodes within a nuclei cluster (as previously defined using the leiden algorithm), was created. TrusTEr(*47*) (dx.doi.org/10.5281/zenodo.7589548) was used to create sample and cluster-specific fastq files. For the processing of the clusters (truster::process_clusters), to retain sample-specific clusters, we defined the grouping parameter to be one per sample (truster::process_clusters, groups). Specific considerations per step in the trusTEr pipeline were taken for the analysis of this dataset:

1. Tsv_to_bam: Due to the large sample size, Rustody’s multi_subset_bam (version 1.3; https://github.com/stela2502/Rustody) was used to subset the individual samples cellranger’s output bam file (-t CB) to individual bam files per sample, per cluster.
2. Filter_UMIs: Samtools view (version 1.18)(*62*) was used to filter out PCR duplicates (as labelled by cellranger).
3. Bam_to_fastq: 10x Genomics bamtofastq (version 1.4.1) was used to retrieve fastq files from the pseudobulk bam files from step 1.
4. Unique mapping: To map reads uniquely to the genome, the flag “unique” was set to True upon execution of the function process_clusters(). STAR aligner (version 2.7.8a; REF; RFID)(*62*) was used using hg38 as the reference genome (--genomeDir), and gencode annotation version 38 as a guide (--sjdbGTFfile). Reads mapping to more than one genomic location (--outFilterMultimapNmax 1) or with a higher mismatch ratio of 0.03 (--outFilterMismatchNoverLmax 0.03) were discarded.
5. TE quantification: Uniquely mapped reads were used to quantify TE expression using featureCounts (subread package, version 1.6.3; REF; RFID, -s 1)(*63*) and repeatmasker (open-4.0.5, filtered from tRNAs, simple repeats, small RNAs, and low-complexity regions) or retrotector predictions for hg38 as annotations.
6. Gene quantification: For methodological consistency and to avoid false discoveries as a result of pseudoreplication, common in single-cell tools for gene differential expression analysis(*64*), we quantified genes also using trusTEr’s pseudobulks (include_genes = True). Gene expression was quantified using featureCounts (subread package, version 1.6.3; REF; RFID, -s 1)(*63*) and gencode annotation version 38 as annotation.

Further documentation of the pipeline can be found at https://molecular-neurogenetics.github.io/truster/.

To visualize the signal coming from each nucleus within the pseudobulk using the same mapping strategy (e.g., Figure S3A), individual tsv files (one per nucleus) were provided as input to trusTEr at step 1, and steps 1 to 5 were executed separately for each nucleus.

#### Gene differential expression and gene set enrichment analyses

Read counts per pseudobulk were further processed in R (version 4.4.1, set.seet(10)). For the tests shown in figures, the following operations were performed per cluster:

1. Remove genes from count matrices with no counts across all pseudobulks.
2. Given the sparse nature of the data, a pseudocount of 1 was added to all genes and all pseudobulks.
3. For clusters that after this filtering had more than three samples per condition (PD vs Control), gene differential expression analysis was performed using DESeq2 (design = ∼ condition) (version 1.44.0; REF; RFID)(*65*).
4. Gene set enrichment analysis was performed using clusterProfiler (version 4.12.1, REF; RFID). Ranked list of genes based on log2FC were passed to the gseGO() function (ont = ALL, keyType = SYMBOL, minGSSize = 3, maxGSSize = 800, seed = T, pvalueCutoff = 0.05, OrgDb = org.Hs.eg.db (version 3.19.1), pAdjustMethod = BH)(*66*).

#### TE differential expression analysis

Read counts per pseudobulk were further processed in R (version 4.4.1, set.seet(10)). For the tests shown in figures, the following operations were performed per cluster:

1. Remove genes from count matrices with no counts across all pseudobulks.
2. Given the sparse nature of the data, a pseudocount of 1 was added to all TEs and all pseudobulks.
3. For clusters that after this filtering had more than three samples per condition (PD vs Control), gene differential expression analysis was performed using DESeq2 (design = ∼ condition) (version 1.44.0; REF; RFID)(*65*).

#### Validation of differential expression analyses

##### Using averages across single nuclei

To validate our results using a more standard approach, we used cellranger’s count matrices (generated using --include_introns) to calculate fold changes over genes (Supplementary Figure 4A). These values were calculated as:

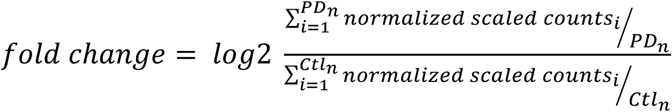

Where PD_n_ is the number of PD nuclei in the cluster, and Ctl_n_ is the number of control nuclei in the cluster.

##### Using other statistical approaches

To assess the robustness of our statistical approach, differential expression analysis using a stringent, more TE-penalizing approach, was used (Figure S3H-J, 4C-D). Read count matrices of the 1259 expressed TEs (as identified from the deep bulk RNAseq) were concatenated with the read count matrix from all genes. The validation run was performed:

1. Removing genes or TEs from the count matrix with no counts across all pseudobulks.
2. Removing pseudobulks with less than 100 nuclei.
3. Adding a pseudocount of 1 to all TEs/genes and all pseudobulks.
4. For clusters that after this filtering had more than three samples per condition (PD vs Control), gene differential expression analysis was performed using DESeq2 (design = ∼ batch + sex + condition) (version 1.44.0; RRID:SCR_015687)(*65*).
5. Gene set enrichment analysis was performed using clusterProfiler (version 4.12.1, RRID:SCR_016884)(*66*). Ranked list of genes based on log2FC were passed to the gseGO() function (ont = ALL, keyType = SYMBOL, minGSSize = 3, maxGSSize = 800, seed = T, pvalueCutoff = 0.05, OrgDb = org.Hs.eg.db (version 3.19.1), pAdjustMethod = BH) (Figure S4D)(*66*).

To evaluate the relevance of covariate adjustment in our data, mean covariate variance across genes/TEs were calculated using a linear mixed model (fitExtractVarPartModel, variancePartition package; RRID:SCR_019204)(*67*) based on transformed counts and precision weights estimated using voom (limma package; RRID:SCR_010943)(*68*) (Table S2).

### Visualization of TE expression using pseudobulks

All heatmaps were produced using pheatmap (version 1.0.12, RRID:SCR_016418). Heatmaps in Figure 2 were plotted using only pseudobulks bigger than 50 nuclei and both heatmaps and boxplots were normalized by the TE group size factors of the pseudobulk (DESeq2 version 1.44.0; RRID:SCR_015687)(*65*). For >6kbp L1HS-PA3 heatmaps (Figure 3 B, D, F, H) and mean plots (Figure 4 B, E), size factors were calculated using only expressed >6kbp L1HS-PA3 counts. For ERV heatmaps (Figure 3 B, D, F, H and Figure 4 C, F), mean plots (Figure 4 B, E), and correlation plots (Figure 6H), size factors were calculated using only expressed ERV counts. All box plots were normalized using gene size factors. To visualize pseudobulks on genome tracks, pseudobulk bam files (see above, step 4 in methods “Pseudobulks”) were indexed using samtools index (version 1.16.1; RRID:SCR_002105)(*69*) and converted to bigwig files using deeptools bamCoverage (version 2.5.4; RRID:SCR_016366)(*70*) using a scale factor of 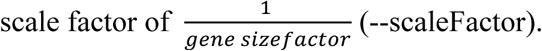.

To visualize the signal from each nucleus within the pseudobulk (e.g., Figure S3A), individual nucleus bam files were indexed using samtools index (version 1.16.1; RRID:SCR_002105) and converted to bigwig files using deeptools bamCoverage (version 2.5.4; RRID:SCR_016366) with RPKM normalization.

### UMAP dimensionality reduction of TE-pseudobulks

ERV or L1 count matrices were normalized by gene size factors of the pseudobulk (DESeq2, (version 1.44.0; RRID:SCR_015687)(*65*) and input to the umap function (umap package, version 0.2.10.0).

### ERV predictions annotation

Genomic coordinates of the ERV predictions were retrieved using Retrotector database version 150418 from elsewhere(*47, 71*). To annotate ERV predictions to the different ERV subfamilies, the coordinates were overlapped with repeatmasker’s annotation and classified based on the largest element (internal regions) overlapping the region, following subfamilies’ classification described elsewhere(*72*). If this was not clear, no internal region was annotated by repeatmasker, or two or more ERV subfamilies were present in the region, the largest or most abundant LTR or internal region fragments overlapping the region were used to classify the prediction.

#### Deep bulk RNA sequencing from post-mortem human brain tissue

A 3 mm^3^ tissue piece was cut from the same tissue block that had been used for snRNA sequencing and disrupted using a Tissuelyser (Qiagen). A steel bead and RLT buffer with β-mercaptoethanol was added to the Tissuelyser and shaken at 30 Hz for 2 minutes. Total RNA was isolated from the disrupted tissue using the RNeasy Mini Kit (Qiagen). The sequencing libraries were then generated using Illumina TruSeq Stranded mRNA library prep kit (with poly-A selection) and sequenced on a NovaSeq6000 or Novaseq X plus (2 x 150 paired end).

#### Bioinformatic analysis of deep bulk RNA sequencing

Reads were uniquely mapped to hg38 reference genome using STAR aligner (version 2.7.8a; RRID:SCR_004463, --outFilterMultimapNmax 1, --outFilterMismatchNoverLmax 0.03)(*62*). TE and gene quantification was performed using featureCounts (subread package version 1.6.3; RRID:SCR_012919; -s 2)(*63*) and gencode annotation version 38, repeatmasker (open-4.0.5, filtered from tRNAs, simple repeats, small RNAs, and low-complexity regions), or retrotector predictions annotation.

Gene size factors were calculated using DESeq2 (version 1.44.0; RRID:SCR_015687)(*65*) and used as a normalization factor for TE expression. Elements were considered to be expressed if they scored over 10 reads in a third of the samples of any group, a group being a cluster of a particular region in one condition (e.g. element X was considered to be expressed, as it had > 10 reads in more than a third of the microglia PUT PD samples).

### ONT direct RNA sequencing

RNA extraction and Poly(A) mRNA isolation were performed as described elsewhere(*46*). The library for long-read direct RNA sequencing was prepared according to manufacturer’s protocol, using the Direct RNA Sequencing Kit (SQK-RNA004) from Oxford Nanopore Technologies. The samples were sequenced using a PromethION 2 Solo IT on a PromethION RNA Flow Cell (FLO-PRO004RA). A detailed protocol can be found at dx.doi.org/10.17504/protocols.io.36wgqpeo3vk5/v1.

### Bioinformatic analysis of ONT direct RNA sequencing

Basecalling was performed using dorado (version 0.7.1) with model rna004_130bps_sup@v3.0.1 and --modified-bases-models using dorado’s model rna004_130bps_sup@v3.0.1_m6A_DRACH@v1. Reads were mapped to the reference genome (hg38) using minimap2 (-ax splice -uf -k14 -y). Alignment files were sorted and indexed using samtools (version 1.16.1). All genome browser tracks were extracted from IgV (version 2.18.2 or 2.15.4; RRID:SCR_011793; http://www.broadinstitute.org/igv/).

### Differentiation and culturing of human microglia from induced pluripotent stem cells

hES (H9; RRID:CVCL_9773) and hiPSCs (KOLF2.1; RRID:CVCL_B5P3) were maintained in IPS-brew on laminin 521 (0.5 μg/cm2) coated plates. The quality control experiments performed for all used iPSC lines are reported at https://doi.org/10.5281/zenodo.17416103. Cells were passaged every 7 days with 0.5 mM EDTA, followed by seeding at a density of 2,500 cells per cm2 with ROCK inhibitor (10 μM Y-27632) included in the medium for the first 24 h after plating. 3×106 iPSCs or ES cells were seeded into an Aggrewell 800 well (STEMCELLTechnologies) to form EBs with ROCK inhibitor (10 μM Y-27632), in iPS-Brew medium and fed daily with medium containing BMP4 (50 ng/ml), VEGF (50 ng/ml), SCF (20 ng/ml). On day 2 and 4 the medium was gently changed in the Aggrewell 800 well plate without agitating the EBs. Day four EBs were then differentiated in either 6-well plates (15 EBs/well), T75 (75 EBs), or T175 flasks (150 EBs) with X-VIVO15 Supplemented with M-CSF (100 ng/ml), IL-3 (25 ng/ml), 2 mM GlutaMAX, 100 U/mL penicillin, 100 mg/mL streptomycin, 0.055 mM b-mercaptoethanol with fresh medium added weekly. pMacpre (Stem cell-derived macrophage/microglia progenitors) emerging into the supernatant after approximately 1 month were collected weekly and differentiation cultures were replenished with fresh medium. Harvested cells were strained (40 mm, Corning) and used: either directly as pMacpre; or plated onto tissue-culture treated plastic or glass coverslips at 100,000 per cm2 and differentiated for 7 days or more. For serum-free differentiation, cells in suspension were harvested and cultured in 75% IMDM, 25% RPMI medium containing B-27 supplement (1:500), l-glutamine (1:1000) and IL-34 (100 ng/ml) and M-CSF (20 ng/ml) for 11–14 days. A detailed protocol can be found at doi: dx.doi.org/10.17504/protocols.io.14egnr8ezl5d/v1.

### Differentiation and culture of hPSC-derived dopaminergic neurons

Human embryonic stem cells (RC17, hPSCreg RCe021-A; RRID:CVCL_L206) were maintained in IPS-brew on laminin 521 (0.5 μg/cm2) coated plates. Cells were passaged every 5 days with 0.5 mM EDTA, followed by seeding at a density of 2,500 cells per cm2 with ROCK inhibitor (10 μM Y-27632-only first 24 h after plating). RC17 ES cells were differentiated towards ventral midbrain (vMB) fate based on a previously published protocol (Nolbrant et al, 2017)(*40*). Day 16 progenitors were plated for terminal differentiation on Lam-521 (2 μg/cm2) plates in Neurobasal medium containing NB-21 supplement without vitamin A (1:500), penicillin/streptomycin (1:1000), l-glutamine (1:1000), BDNF (20 ng/ml), AA (0.2 mM), GDNF (10 ng/ml), db-cAMP (500 μM) and DAPT (1 μM). Detailed protocols for generation of human ventral midbrain dopaminergic progenitors and dopaminergic neurons can be found at dx.doi.org/10.17504/protocols.io.kxygx4eqol8j/v1 and dx.doi.org/10.17504/protocols.io.q26g7nbd1lwz/v1, respectively.

### Immunocytochemistry

For immunocytochemistry (ICC), cells were washed with 1× phosphate-buffered saline (DPBS) and fixed with 4% paraformaldehyde (PFA) in DPBS for 10 minutes at room temperature (RT). Following fixation, cells were washed three times with DPBS and stored at 4 °C until further use.

Cells were incubated in blocking solution (5% donkey serum and 0.1% Triton X-100 in DPBS) for 1 hour at RT, followed by overnight incubation at 4 °C with primary antibodies (Table 1) diluted in the same blocking solution. The next day, cells were washed twice with DPBS and incubated with fluorophore-conjugated secondary antibodies in blocking solution for 1 hour at RT. Afterwards, cells were washed twice with DPBS and counterstained with DAPI (D9542, Sigma-Aldrich) diluted in DPBS for 5 minutes at RT. Finally, cells were washed an additional two times with DPBS before imaging.

**Table 1.** Primary antibodies used for immunocytochemistry. The table includes target proteins, catalogue numbers and vendors. Antibodies were validated according to the manufacturers’ recommendations and used under optimised experimental conditions.

| Name | Cat.no | Vendor |
| --- | --- | --- |
| CD45 | 304001 | Biolegend |
| IBA1 | ab5076 | Abcam |
| PU1 | 2258S | Cell Signaling |
| TREM2 | MAB17291 | R&D |

Immunofluorescent images were acquired at 20X with a confocal microscope (CV8000, Yokogawa). A detailed protocol can be found at doi: dx.doi.org/10.17504/protocols.io.5jyl88m69l2w/v1. Immunocytochemistry images included in the paper are deposited on Zenodo at the following link: https://doi.org/10.5281/zenodo.17405239.

### Stimulation of DA neurons and microglia

Terminally differentiated DA neurons and microglia were stimulated with IFNg (5ng/ml), and TNFa (5ng/ml) for 48hrs in the culture medium. IFNg and TNFa were reconstituted in PBS. Control cells were treated with PBS only. Post stimulation, cells were washed twice with 1XPBS and detached with Accutase and pellets were frozen down on dry ice for bulk-RNA seq. A detailed protocol can be found at doi: dx.doi.org/10.17504/protocols.io.261geko2dg47/v1.

#### Bioinformatic analysis of stimulated DA neurons and microglia

Reads were uniquely mapped to hg38 reference genome using STAR aligner (version 2.7.8a; RRID:SCR_004463, --outFilterMultimapNmax 1, --outFilterMismatchNoverLmax 0.03)(*62*). TE and gene quantification was performed using featureCounts (subread package version 1.6.3; RRID:SCR_012919; -s 2)(*63*) and gencode annotation version 38, repeatmasker (open-4.0.5, filtered from tRNAs, simple repeats, small RNAs, and low-complexity regions), or retrotector predictions annotation.

Gene differential expression analysis was performed on DA neurons or microglia and for each of the treatments separately using DESeq2 (design = ∼ treatment) (version 1.44.0; RRID:SCR_015687)(*65*). Gene set enrichment analysis was performed using clusterProfiler (version 4.12.1, RRID:SCR_016884)(*66*). Ranked list of genes based on log2FC were passed to the gseGO() function (ont = ALL, keyType = SYMBOL, minGSSize = 3, maxGSSize = 800, seed = T, pvalueCutoff = 0.05, OrgDb = org.Hs.eg.db (version 3.19.1), pAdjustMethod = BH). TE differential expression analysis was performed on DA neurons or microglia and for each of the treatments separately using DESeq2 (design = ∼ treatment) (version 1.44.0; RRID:SCR_015687)(*65*). Gene size factors were calculated using DESeq2 and used as a normalization factor for TE expression visualization (Figure 7J).

## ACKNOWLEDGMENTS

This research was funded by Aligning Science Across Parkinson’s (Grant IDs: ASAP-000520, ASAP-024296 and ASAP-025170) through the Michael J. Fox Foundation for Parkinson’s Research (MJFF). J.J received support from the Swedish Government Initiative for Strategic Research Areas (MultiPark & StemTherapy). A.K. received support by the Novo Nordisk Foundation (NNF21CC0073729) and by Bioneer A/S (DigitStem grant). J.Jones., A.Q and R.A.B. received support from the NIHR Cambridge Biomedical Research Centre (BRC-1215-20014). The Cambridge Brain Bank is supported by the NIHR Cambridge Biomedical Research Centre (NIHR203312). The views expressed here are those of the authors, and not necessarily of the NIHR, Department of Health and Social Care. The funding bodies played no role in the design of the study and collection, analysis, and interpretation of data and in writing the manuscript.

## Funding

### Author contributions

Conceptualization: R.G., A.A., A.T., A.C., M.H., A.K., R.B., J. Jakobsson.

Methodology: R.G., A.A., A.T., V.H., J. Johansson, A.K., R.B.

Software: R.G., T.F., Y.S.

Validation: R.G., A.A., A.T., A.C., T.F., D.A.

Formal analysis: R.G., A.T., O.T., Y.S.

Investigation: A.A., A.T., D.L., N.K., D.A., V.H., S.B., J. Johansson, D.R., J. Jones, A.Q., R.B.

Resources: A.T., S.W., A.C., Y.S., A.Q., R.B.

Data curation: R.G., A.C., O.T., D.R., L.C., A.Q., M.H.

Writing - original draft: R.G., A.A., A.T., R.B., J. Jakobsson.

Writing - review and editing: R.G., A.A., A.T., S.W., A.C., O.T., T.F., N.K., Y.S., D.R., L.C., J. Jones, A.Q., M.H., A.K., R.B., J. Jakobsson.

Visualization: R.G., A.T., N.K., D.A.

Supervision: R.G., J. Jones, A.Q., M.H., A.K., R.B., J. Jakobsson.

Project administration: R.G., A.A., L.C., A.Q., A.K., R.B., J. Jakobsson.

Funding acquisition: M.H., A.K., R.B., J. Jakobsson.

### Data, Code, and Materials Availability

This study did not generate new materials. The RC17 cell line can be provided by ROSLIN CELLS LIMITED pending scientific review and execution of a completed material transfer agreement. The KOLF2.1 cell line can be provided by The Jackson Laboratory pending scientific review and execution of a completed material transfer agreement. The H9 cell line can be provided by WiCell Research Institute pending scientific review and execution of a completed material transfer agreement. Human material can be provided by the Cambridge Brain Bank pending scientific review and execution of a completed material transfer agreement. Requests should be directed to the corresponding authors, who will facilitate contact with the appropriate providing institution. All the data, code (https://doi.org/10.5281/zenodo.17434560) and protocols generated and used in this study needed to evaluate and reproduce the results in the paper have been deposited in public repositories. They are listed, together with used key lab materials, alongside their persistent identifiers in a Key Resource Table at https://doi.org/10.5281/zenodo.18863546. Data used in the preparation of this article was obtained from the following collections from the Aligning Science Across Parkinson’s Collaborative Research Network Cloud (ASAP CRN Cloud) (RRID:SCR_023923)(*73*): “Single nuclei sequencing of brain regions from healthy and Parkinson’s Disease individuals” (https://doi.org/10.5281/zenodo.15162834), “Deep bulk RNAseq of neurological controls and PD brains” (https://doi.org/10.5281/zenodo.16929448), “Bulk RNAseq of dopaminergic neurons in vitro cultures” (https://doi.org/10.5281/zenodo.17149267), and “Bulk RNAseq of microglia in vitro cultures” (https://doi.org/10.5281/zenodo.17149291). The data are controlled. Researchers can register for access to these data by submitting a Data Use Application through the ASAP CRN Cloud website (https://cloud.parkinsonsroadmap.org/collections). Data dictionaries, README files, protocols used to collect the data, and data processing pipelines are openly available at https://cloud.parkinsonsroadmap.org/collections.

We want to thank Patric Jern for his support throughout this project and M. Persson Vejgården, A. Hammarberg and U. Jarl for their technical assistance. We are grateful to all members of the Jakobsson lab. The views expressed here are those of the authors, and not necessarily of the NIHR, Department of Health and Social Care.

## TABLE OF CONTENTS FOR SUPPLEMENTAL MATERIALS

**Figure S1:** snRNA-seq quality control metrics, canonical markers gene expression, comparison of cell type composition between PD and control, and condition-specific UMAP visualizations.

**Figure S2:** Deep bulk RNA-seq quality control graphs.

**Figure S3:** Validation analyses of cell-type specific TE expression from snRNA-seq.

**Figure S4:** Classification of ERVs upregulated in PD microglia (SN and PUT), differential expression analysis (DEA) of ERVs and L1s in the remaining cell types, and validation of TE DEA results with covariate correction.

**Figure S5:** Validation of gene DEA results using group means, and covariate correction.

**Figure S6:** Validation of in vitro findings using hiPSC-derived microglia from an independent cell line (KOLF).

**Table S1:** Datasets and donors’ information for the post-mortem cohort.

**Table S2:** Mean variance explained by individual covariates.

