## Supplemental Figures and Tables for "Activation of transposable elements is linked to a region- and cell-type-specific interferon response in Parkinson’s disease"

### SUPPLEMENTAL MATERIAL

#### *Supplemental Figures*

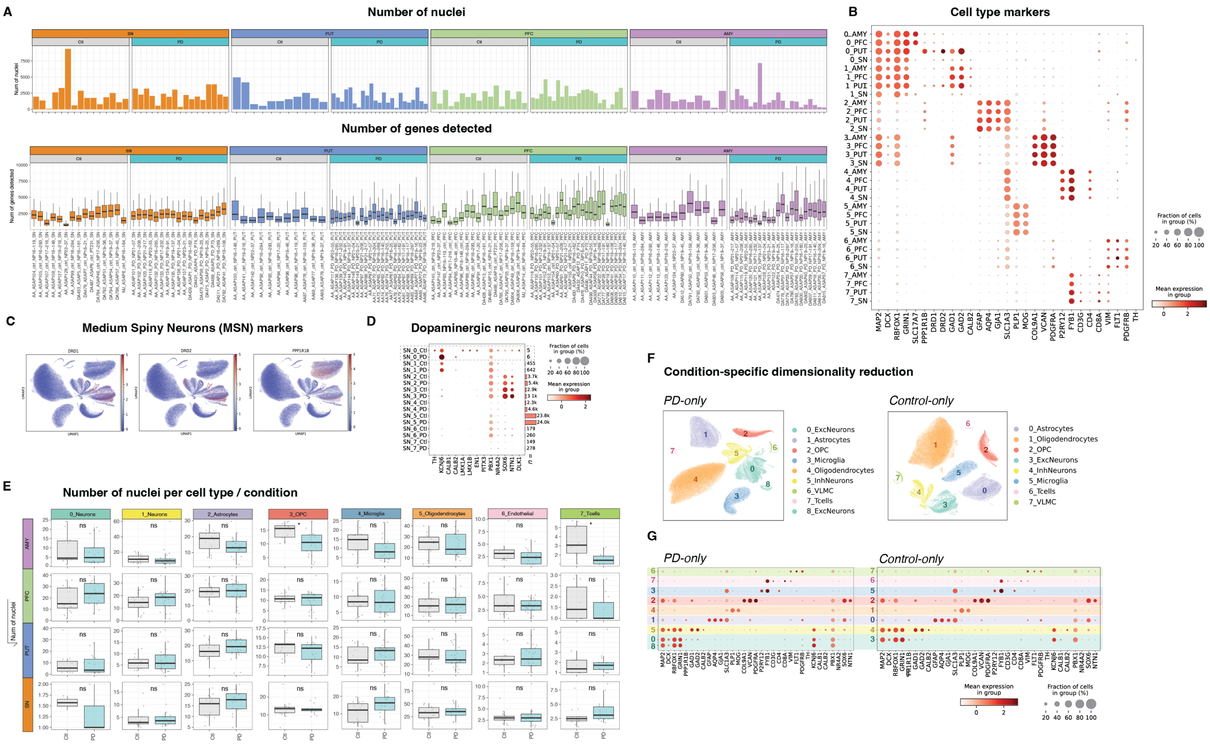


***Figure S1. snRNA-seq quality control metrics, canonical markers gene expression, comparison of cell type composition between PD and control, and condition-specific UMAP visualizations. A)*** *Number of nuclei (top) and genes detected (bottom) in each sample.* ***B)*** *Dotplot showing canonical gene markers in each cluster and region. Dot size corresponding to the fraction of nuclei in cluster expressing the marker. Color representing average gene expression in cluster.* ***C)*** *UMAPs coloured by normalized expression of MSN gene markers.* ***D)*** *Dotplot showing canonical dopaminergic neuron markers in each cluster in the SN. Dot size corresponding to the fraction of nuclei in cluster expressing the marker. Color representing average gene expression in cluster.* ***E)*** *Boxplot showing the number of nuclei per individual in each cluster (y-axis: squared root of the number of nuclei per sample) grouped by condition. T-tests showing a statistically significant difference in the number of nuclei in each cluster.* ***F)*** *UMAP showing unbiased clustering of all nuclei from PD (left; 9 clusters) and Control (right; 8 clusters) individuals. Clusters are colored by cell type.* ***G)*** *Dotplot showing canonical gene markers expression from PD (left) and Control (right) clusters. Dot size corresponding to the fraction of nuclei in cluster expressing the marker. Color representing average gene expression in cluster.*

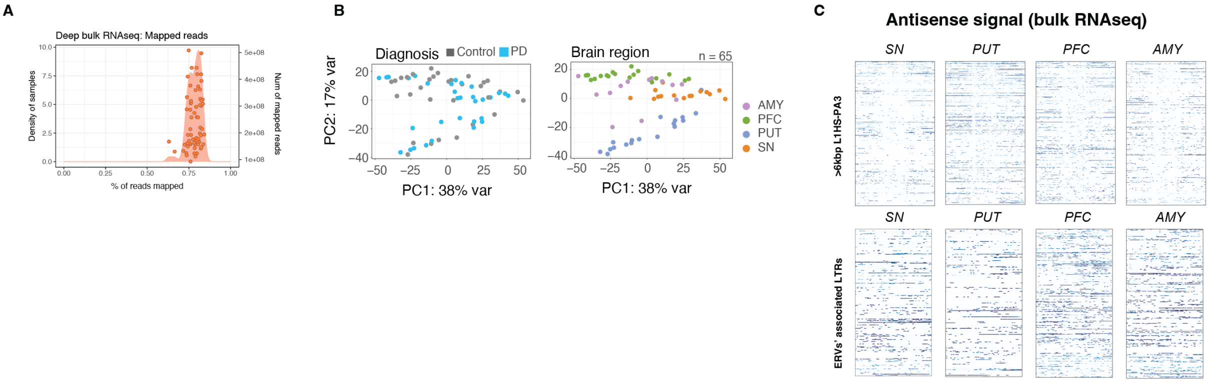


***Figure S2. Deep bulk RNA-seq quality control graphs. A)*** *Distribution of number of reads and percentage of reads mapped to the genome among the deep bulk RNA-seq samples.* ***B)*** *Principal component analysis of the samples sequenced with deep bulk RNA-seq plotted by diagnosis (top: Diagnosis) or brain region (bottom: Brain region). AMY= amygdala; PFC= prefrontal cortex; PUT= putamen; SN= substantia nigra* ***C)*** *Top: Antisense expression heatmap showing binned signal (10bp) over expressed >6kbp L1HS-L1PA3 elements (from Figure 2b, n = 520) with 6kbp windows up and downstream. Elements are scaled to 6kbp. Color represents log2 RPKM values. Bottom: Antisense expression heatmap showing binned signal (10bp) over the associated LTRs to the expressed ERV elements (from Figure 2b, n = 739) with 5kbp windows up and downstream. Elements are scaled to 1kbp. Color represents log2 RPKM values.*

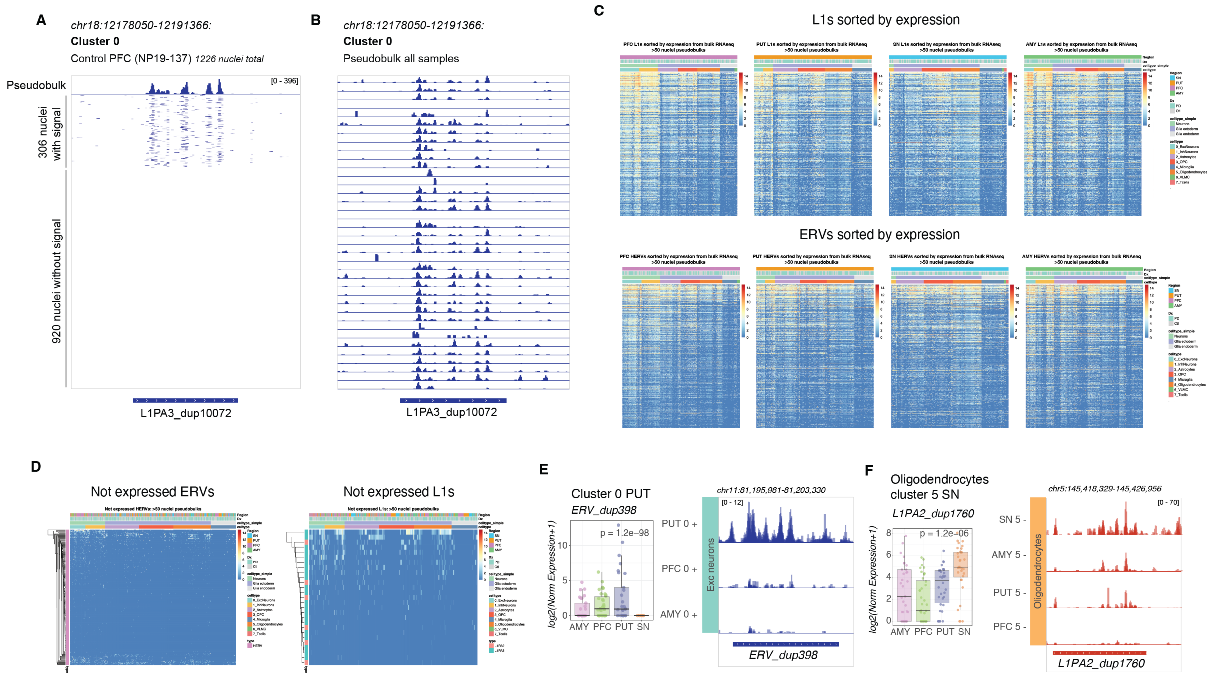


***Figure S3. Validation analyses of cell-type specific TE expression from snRNA-seq. A)*** *Genome browser tracks showing, at the top, the pseudobulk of a single sample (cluster 0 of NP19-137 PFC, normalized by gene size factors), followed by 1226 individual nucleus tracks contributing to the pseudobulk (normalized by RPKM). Positive-strand transcription is shown.* ***B)*** *Genome browser tracks showing all cluster 0 pseudobulks across samples. Tracks were normalized by gene size factors, positive-strand transcription is shown.* ***C)*** *Heatmaps showing cell-type (columns) TE (rows) expression in the different brain regions. ERVs (left) and >6kbp L1HS-L1PA3 (right) shown were found not to be expressed in the deep-bulk RNAseq. Column annotation showing cell-types, condition (Dx, PD or Ctl), and brain region.* ***D)*** *Heatmaps showing cell-type (columns) TE (rows) expression in the different brain regions. ERVs (left) and >6kbp L1HS-L1PA3 (right) ordered by expression in deep-bulk RNAseq from the respective brain region. Column annotation showing cell-types, condition (Dx, PD or Ctl), and brain region.* ***E)*** *Example of an excitatory neuron-, putamen-specific ERV. To the left, boxplot showing samples' cell-type expression (DESeq2, Wald test, normalized by gene size factors). To the right, genome browser tracks showing excitatory neurons in PUT, AMY and PUT (normalized by gene size factors).* ***F)*** *Example of an oligodendrocyte-, SN-specific L1PA2. To the left, boxplot showing cell-type expression (DESeq2, Wald test, normalized by gene size factors). To the right, genome browser tracks showing excitatory neurons in PUT, AMY and PUT (normalized by gene size factors). Genome tracks in blue showing transcription from the positive strand, red showing transcription from the negative strand.*

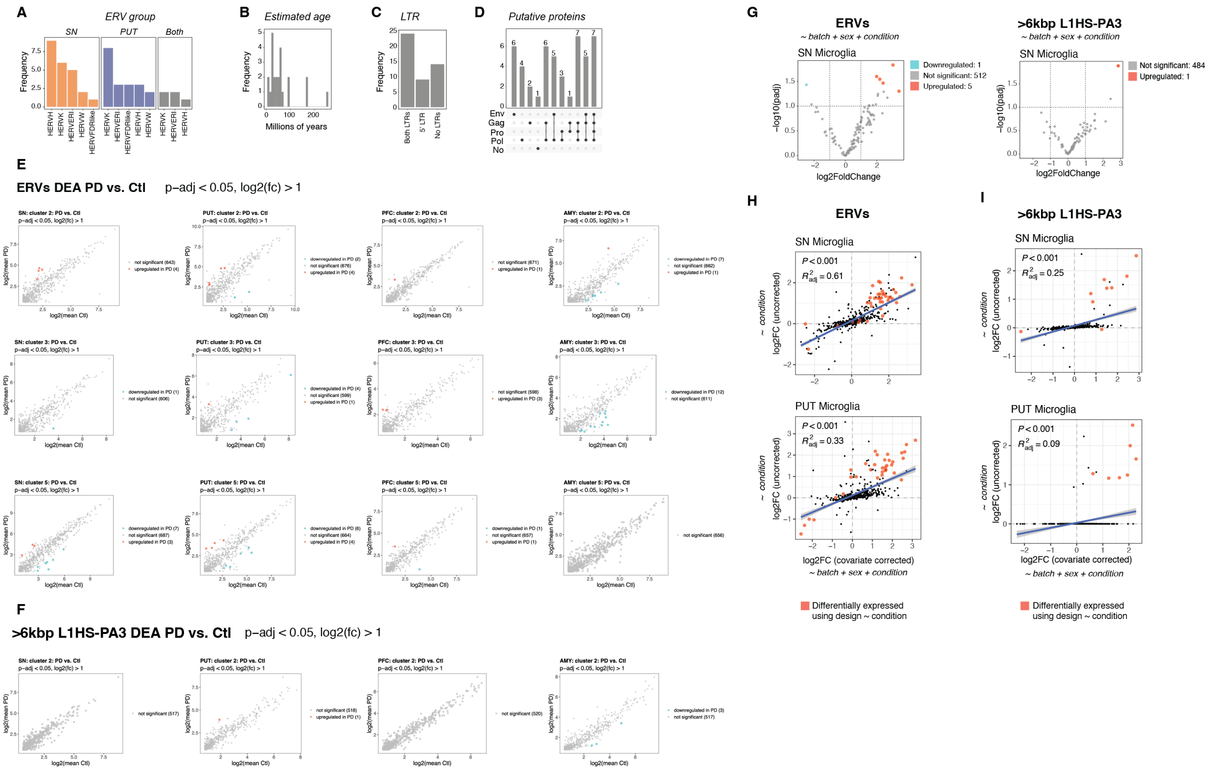


***Figure S4. Classification of ERVs upregulated in PD microglia (SN and PUT), differential expression analysis (DEA) of ERVs and L1s in the remaining cell types, and validation of TE DEA results with covariate correction. A)*** *Subfamilies of upregulated ERVs in SN and PUT microglia.* ***B)*** *Estimated evolutionary age of upregulated ERVs in PD SN and PUT microglia.* ***C)*** *Presence of LTRs in upregulated ERVs in PD SN and PUT microglia.* ***D)*** *Putative proteins as reported by retrotector in upregulated ERVs in PD SN and PUT microglia.* ***E)*** *Mean plots showing ERV differential expression analysis results in the cell types not shown in main figure for each brain region (red indicating TEs of padj < 0.05 & log2FC > 1. DESeq2 Wald-test).* ***F)*** *Mean plots showing >6kbp L1HS-PA3 differential expression analysis results in the cell types not shown in main figure for each brain region (red indicating TEs of padj < 0.05 & log2FC > 1. DESeq2 Wald-test).* ***G)*** *Volcano plot showing ERV (left) and >6kbp L1HS-PA3 (right) DEA in SN microglia using design ~ batch + sex + condition (red indicating TEs of padj < 0.05 & log2FC > 1. DESeq2 Wald-test).* ***H-I)*** *Scatter plots showing correlation between log2FC calculated with (x-axis) and without (y-axis) covariate correction. Red indicates TEs of padj < 0.05 & |log2FC| > 1 using uncorrected test (TE log2FC and padj calculated by DESeq2 Wald-test. Fitted linear models using method = lm, ggpmisc::stat_poly_line).*

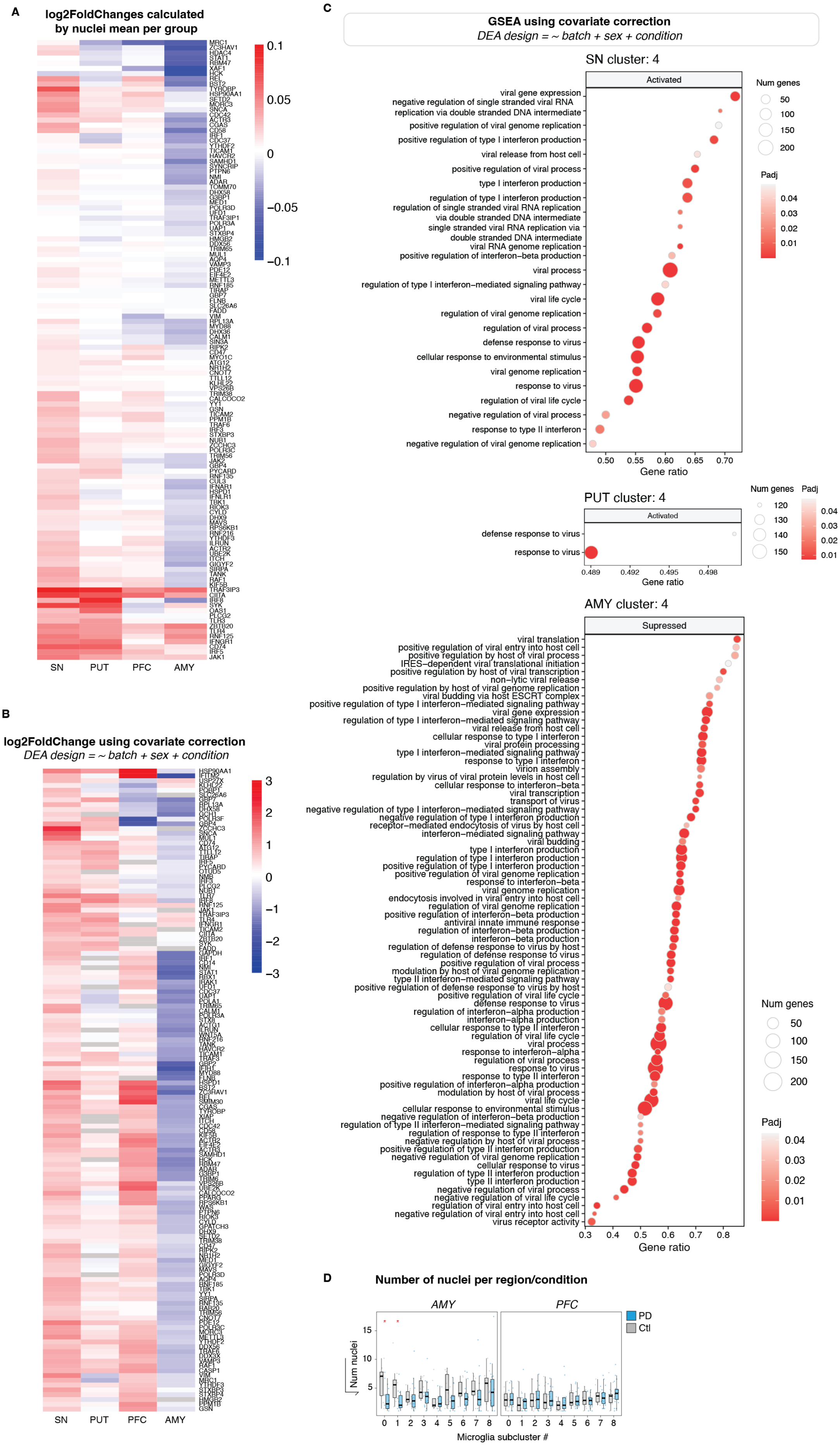
A
log2FoldChanges calculated by nuclei mean per group
SN
PUT
PFC
AMY
B
log2FoldChange using covariate correction
DEA design = ~ batch + sex + condition
C
GSEA using covariate correction
DEA design = ~ batch + sex + condition
SN cluster: 4
Activated
viral gene expression
negative regulation of single stranded viral RNA replication via double stranded DNA intermediate
positive regulation of viral genome replication
positive regulation of type I interferon production
viral release from host cell
positive regulation of viral process
type I interferon production
regulation of type I interferon production
regulation of single stranded viral RNA replication via double stranded DNA intermediate
single stranded viral RNA replication via double stranded DNA intermediate
viral RNA genome replication
positive regulation of interferon−beta production
viral process
regulation of type I interferon−mediated signaling pathway
viral life cycle
regulation of viral genome replication
regulation of viral process
defense response to virus
cellular response to environmental stimulus
viral genome replication
response to virus
regulation of viral life cycle
negative regulation of viral process
response to type II interferon
negative regulation of viral genome replication
Gene ratio
Num genes
Padj
PUT cluster: 4
Activated
defense response to virus
response to virus
Gene ratio
AMY cluster: 4
Supressed
viral translation
positive regulation of viral entry into host cell
positive regulation by host of viral process
IRES−dependent viral translational initiation
positive regulation by host of viral transcription
non−lytic viral release
positive regulation by host of viral genome replication
viral budding via host ESCRT complex
positive regulation of type I interferon−mediated signaling pathway
viral gene expression
regulation of type I interferon−mediated signaling pathway
viral release from host cell
cellular response to type I interferon
viral protein processing
type I interferon−mediated signaling pathway
response to type I interferon
virion assembly
regulation by virus of viral protein levels in host cell
cellular response to interferon−beta
viral transcription
transport of virus
negative regulation of type I interferon−mediated signaling pathway
negative regulation of type I interferon production
receptor−mediated endocytosis of virus by host cell
interferon−mediated signaling pathway
viral budding
type I interferon production
regulation of type I interferon production
positive regulation of type I interferon production
positive regulation of viral genome replication
response to interferon−beta
viral genome replication
endocytosis involved in viral entry into host cell
regulation of viral genome replication
positive regulation of interferon−beta production
antiviral innate immune response
regulation of interferon−beta production
interferon−beta production
regulation of defense response to virus by host
regulation of defense response to virus
positive regulation of viral process
modulation by host of viral genome replication
type II interferon−mediated signaling pathway
positive regulation of defense response to virus by host
positive regulation of viral life cycle
defense response to virus
regulation of interferon−alpha production
interferon−alpha production
cellular response to type II interferon
regulation of viral life cycle
viral process
response to interferon−alpha
regulation of viral process
response to virus
response to type II interferon
positive regulation of interferon−alpha production
modulation by host of viral process
viral life cycle
cellular response to environmental stimulus
negative regulation of interferon−beta production
regulation of type II interferon−mediated signaling pathway
regulation of response to type II interferon
negative regulation by host of viral process
positive regulation of type II interferon production
negative regulation of viral genome replication
cellular response to virus
regulation of type II interferon production
type II interferon production
negative regulation of viral process
negative regulation of viral life cycle
regulation of viral entry into host cell
negative regulation of viral entry into host cell
virus receptor activity
Gene ratio
Num genes
Padj
D
Number of nuclei per region/condition
AMY
PFC
PD
Ctl
Num nuclei
Microglia subcluster #

***Figure S5. Validation of gene DEA results using group means, and covariate correction. A)** Heatmap coloured by stable log2FC (scaled mean in PD vs scaled mean in Control; see methods "Validation of differential expression analyses") of the genes identified as the core enrichment of the GO terms shown in Figure 6. **B)** Heatmap of genes identified as the core enrichment of the GO terms shown in Figure 6 coloured by log2FC in microglia using covariate corrected data (DESeq2, design = ~ batch + sex + condition). **C)** Interferon and viral related GO terms selected from the results of a gene set enrichment analysis (GSEA) of SN (left), PUT (middle), and AMY (right) in PD Microglia (clusterProfiler::gseGO) using covariate correction (DESeq2, design = ~ batch + sex + condition). **D)** Boxplot showing the number of nuclei per individual in each microglia subcluster (y-axis: squared root of the number of nuclei). T-tests showing a statistically significant difference in the number of microglia in AMY (clusters 0 and 1).*

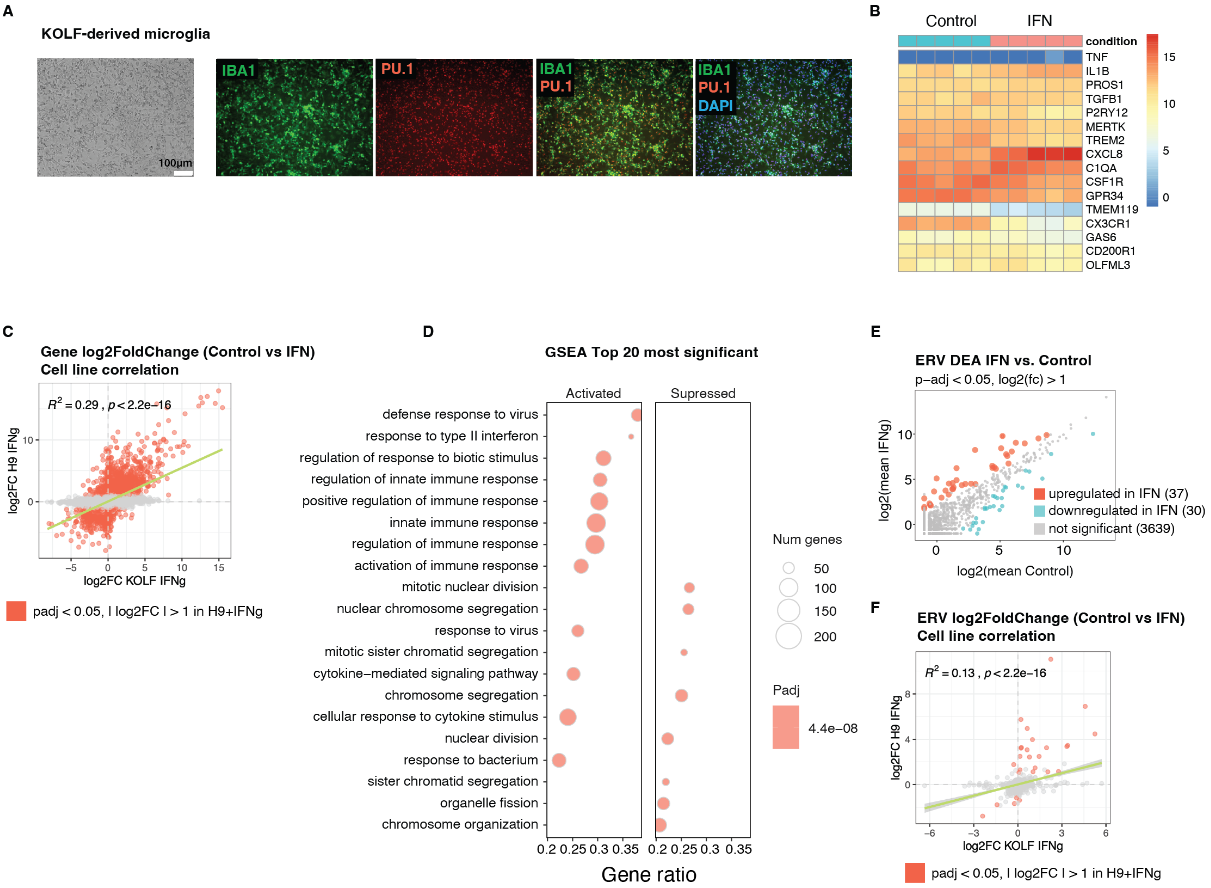


***Figure S6. Validation of in vitro findings using hiPSC-derived microglia from an independent cell line (KOLF). A)** Left: Brightfield image showing KOLF hiPSC-derived microglia at 10 days of in vitro differentiation. Scale bar = 100µm. Right: Immunostaining of microglial markers (PU.1, and IBA-1). Scale bar = 100µm. **B)** Heatmap showing microglial marker expression from bulk RNA sequencing. **C)** Scatter plots showing correlation between genes' log2FC (Untreated vs IFNg treated microglia) in H9-derived microglia (y-axis) and KOLF-derived microglia (x-axis). Red indicates differentially expressed genes in H9-derived microglia (padj < 0.05 & | log2FC | > 1; DESeq2 Wald-test. Fitted linear model using method = lm, ggpmisc::stat_poly_line). **D)** Gene set enrichment analysis (GSEA) results of Untreated vs IFN treated KOLF-derived microglia. **E)** Mean plot showing ERV differential expression analysis results in KOLF-derived microglia (red indicating ERVs of pdj < 0.05 & log2FC >*

*1. DESeq2 Wald-test).* ***F)*** *Scatter plots showing correlation between ERVs log2FC (Untreated vs IFNg treated microglia) in H9-derived microglia (y-axis) and KOLF-derived microglia (x-axis). Red indicates differentially expressed ERVs in H9-derived microglia (padj < 0.05 & | log2FC | > 1; DESeq2 Wald-test. Fitted linear model using method = lm, ggpmisc::stat_poly_line).*

#### *Supplemental Tables*

***Table S1. Datasets and donors' information for the post-mortem cohort.*** *Datasets and donors' information from the 46 individuals within the post-mortem cohort. Information includes the condition (control or Parkinson's disease), brain region (PFC= prefrontal cortex; PUT= putamen; SN= substantia nigra; AMY= amygdala), age, sex (M= male; F= female), post-mortem interval (PMI) and Braak stage.*

| Sample ID | Diagnosis | Region | Age | Sex | PMI | Braak | 10x snRNAseq | Deep Bulk RNAseq |
|---|---|---|---|---|---|---|---|---|
| NP16-00119 | Control | PFC | 74 | M | 74 | 0 | Yes | Yes |
| NP16-00119 | Control | PUT | 74 | M | 74 | 0 | Yes | Yes |
| NP16-00119 | Control | SN | 74 | M | 74 | 0 | Yes | Yes |
| NP16-00119 | Control | AMY | 74 | M | 32 | 0 | Yes | Yes |
| NP16-00284 | Control | PFC | 70 | F | 32 | 0 | Yes | Yes |
| NP16-00284 | Control | PUT | 70 | F | 32 | 0 | Yes | Yes |
| NP16-00284 | Control | SN | 70 | F | 32 | 0 | Yes | Yes |
| NP16-00284 | Control | AMY | 70 | F | 32 | 0 | Yes | Yes |
| NP16-00285 | PD | AMY | 87 | M | 68 | 5 | Yes | Yes |
| NP16-00285 | PD | SN | 87 | M | 68 | 5 | Yes | Yes |
| NP16-00285 | PD | PUT | 87 | M | 68 | 5 | Yes | Yes |
| NP16-00285 | PD | PFC | 87 | M | 68 | 5 | Yes | Yes |
| NP16-00293 | Control | PFC | 75 | F | 92 | 0 | Yes | Yes |
| NP16-00293 | Control | PUT | 75 | F | 92 | 0 | Yes | Yes |
| NP16-00293 | Control | SN | 75 | F | 92 | 0 | Yes | Yes |
| NP16-00293 | Control | AMY | 75 | F | 32 | 0 | Yes | Yes |
| NP16-140 | PD | AMY | 71 | M | 32 | 6 | Yes | |
| NP16-140 | PD | PFC | 71 | M | 32 | 6 | Yes | |
| NP16-140 | PD | PUT | 71 | M | 32 | 6 | Yes | |
| NP16-160 | PD | PFC | 88 | M | 32 | 6 | Yes | |
| NP16-160 | PD | AMY | 88 | M | 32 | 6 | Yes | |
| NP16-160 | PD | PUT | 88 | M | 32 | 6 | Yes | |
| NP16-161 | Control | PFC | 72 | M | 32 | 0 | Yes | |
| NP16-161 | Control | SN | 72 | M | 32 | 0 | Yes | |
| NP16-162 | PD | PUT | 74 | M | 32 | 6 | Yes | |
| NP16-162 | PD | SN | 74 | M | 32 | 6 | Yes | |
| NP16-162 | PD | PFC | 74 | M | 32 | 6 | Yes | |
| NP16-164 | Control | PFC | 75 | M | 32 | 0 | Yes | |
| NP16-164 | Control | SN | 75 | M | 32 | 0 | Yes | |
| NP16-21 | Control | SN | 83 | M | 32 | 0 | Yes | |

| | | | | | | | | |
|---|---|---|---|---|---|---|---|---|
| NP16-21 | Control | PFC | 83 | M | 32 | 0 | Yes | |
| NP16-25 | PD | PUT | 85 | M | 32 | 4 | Yes | |
| NP16-25 | PD | AMY | 85 | M | 32 | 4 | Yes | |
| NP16-25 | PD | SN | 85 | M | 32 | 4 | Yes | |
| NP16-25 | PD | PFC | 85 | M | 32 | 4 | Yes | |
| NP16-269 | PD | PFC | 73 | M | 32 | 5 | Yes | |
| NP16-269 | PD | PUT | 73 | M | 32 | 5 | Yes | |
| NP16-269 | PD | SN | 73 | M | 32 | 5 | Yes | |
| NP17-00020 | Control | PFC | 87 | M | 49 | 0 | Yes | Yes |
| NP17-00020 | Control | PUT | 87 | M | 49 | 0 | Yes | Yes |
| NP17-00020 | Control | AMY | 87 | M | 32 | 0 | | Yes |
| NP17-00216 | Control | PFC | 77 | F | 59 | 0 | Yes | Yes |
| NP17-00216 | Control | PUT | 77 | F | 59 | 0 | | Yes |
| NP17-00216 | Control | SN | 77 | F | 32 | 0 | Yes | Yes |
| NP17-00232 | PD | AMY | 71 | F | 54 | 4 | Yes | Yes |
| NP17-00232 | PD | SN | 71 | F | 54 | 4 | Yes | Yes |
| NP17-00232 | PD | PUT | 71 | F | 54 | 4 | Yes | Yes |
| NP17-00232 | PD | PFC | 71 | F | 54 | 4 | Yes | Yes |
| NP17-191 | PD | PFC | 73 | M | 32 | 6 | Yes | |
| NP17-191 | PD | AMY | 73 | M | 32 | 6 | Yes | |
| NP17-191 | PD | PUT | 73 | M | 32 | 6 | Yes | |
| NP17-256 | Control | SN | 72 | F | 32 | 0 | Yes | |
| NP17-256 | Control | PFC | 72 | F | 32 | 0 | Yes | |
| NP17-256 | Control | AMY | 72 | F | 32 | 0 | Yes | |
| NP18-00046 | Control | PFC | 86 | F | 67 | 0 | Yes | Yes |
| NP18-00046 | Control | PUT | 86 | F | 32 | 0 | Yes | Yes |
| NP18-00148 | Control | PFC | 88 | F | 28 | 0 | Yes | Yes |
| NP18-00148 | Control | PUT | 88 | F | 28 | 0 | Yes | Yes |
| NP18-00148 | Control | SN | 88 | F | 28 | 0 | Yes | Yes |
| NP18-00148 | Control | AMY | 88 | F | 28 | 0 | Yes | Yes |
| NP18-117 | PD | AMY | 90 | M | 32 | 5 | Yes | |
| NP18-117 | PD | PFC | 90 | M | 32 | 5 | Yes | |
| NP18-117 | PD | PUT | 90 | M | 32 | 5 | Yes | |
| NP18-159 | Control | AMY | 65 | F | 32 | 0 | Yes | |
| NP18-159 | Control | PFC | 65 | F | 32 | 0 | Yes | |
| NP18-159 | Control | PUT | 65 | F | 32 | 0 | Yes | |
| NP18-287 | PD | AMY | 77 | M | 32 | 5 | Yes | |
| NP18-287 | PD | PUT | 77 | M | 32 | 5 | Yes | |
| NP18-304 | PD | PFC | 89 | M | 32 | 4 | Yes | |
| NP18-304 | PD | AMY | 89 | M | 32 | 4 | Yes | |
| NP18-304 | PD | PUT | 89 | M | 32 | 4 | Yes | |
| NP19-00091 | PD | AMY | 82 | F | 67 | 5 | Yes | Yes |
| NP19-00091 | PD | SN | 82 | F | 67 | 5 | Yes | Yes |

| | | | | | | | | |
|---|---|---|---|---|---|---|---|---|
| NP19-00091 | PD | PUT | 82 | F | 67 | 5 | Yes | Yes |
| NP19-00091 | PD | PFC | 82 | F | 32 | 5 | Yes | Yes |
| NP19-00137 | PD | AMY | 77 | M | 59 | 5 | Yes | Yes |
| NP19-00137 | PD | SN | 77 | M | 59 | 5 | Yes | Yes |
| NP19-00137 | PD | PUT | 77 | M | 59 | 5 | Yes | Yes |
| NP19-00137 | PD | PFC | 77 | M | 59 | 5 | Yes | Yes |
| NP19-00218 | Control | AMY | 88 | M | 18 | 0 | Yes | Yes |
| NP19-00218 | Control | SN | 88 | M | 18 | 0 | Yes | Yes |
| NP19-00218 | Control | PUT | 88 | M | 18 | 0 | Yes | Yes |
| NP19-00218 | Control | PFC | 88 | M | 18 | 0 | Yes | Yes |
| NP19-108 | PD | SN | 86 | F | 32 | 6 | Yes | |
| NP19-108 | PD | PFC | 86 | F | 32 | 6 | Yes | |
| NP19-108 | PD | PUT | 86 | F | 32 | 6 | Yes | |
| NP19-16 | PD | AMY | 84 | M | 32 | 4 | Yes | |
| NP19-16 | PD | PUT | 84 | M | 32 | 4 | Yes | |
| NP19-16 | PD | PFC | 84 | M | 32 | 4 | Yes | |
| NP19-23 | PD | AMY | 78 | F | 32 | 5 | Yes | |
| NP19-23 | PD | PUT | 78 | F | 32 | 5 | Yes | |
| NP19-23 | PD | SN | 78 | F | 32 | 5 | Yes | |
| NP19-255 | PD | AMY | 76 | M | 32 | 5 | Yes | |
| NP19-255 | PD | PUT | 76 | M | 32 | 5 | Yes | |
| NP19-255 | PD | PFC | 76 | M | 32 | 5 | Yes | |
| NP19-36 | Control | SN | 39 | M | 32 | 0 | Yes | |
| NP19-36 | Control | AMY | 39 | M | 32 | 0 | Yes | |
| NP19-36 | Control | PUT | 39 | M | 32 | 0 | Yes | |
| NP19-37 | Control | SN | 73 | M | 32 | 0 | Yes | |
| NP19-37 | Control | PFC | 73 | M | 32 | 0 | Yes | |
| NP19-37 | Control | AMY | 73 | M | 32 | 0 | Yes | |
| NP19-37 | Control | PUT | 73 | M | 32 | 0 | Yes | |
| NP19-45 | Control | SN | 62 | M | 32 | 0 | Yes | |
| NP19-45 | Control | PFC | 62 | M | 32 | 0 | Yes | |
| NP19-45 | Control | AMY | 62 | M | 32 | 0 | Yes | |
| NP21-00004 | PD | AMY | 75 | M | 48 | 6 | Yes | Yes |
| NP21-00004 | PD | SN | 75 | M | 48 | 6 | Yes | Yes |
| NP21-00004 | PD | PUT | 75 | M | 48 | 6 | Yes | Yes |
| NP21-00004 | PD | PFC | 75 | M | 48 | 6 | Yes | Yes |
| NP21-00057 | PD | PFC | 83 | F | 28 | 4 | Yes | Yes |
| NP21-00057 | PD | PUT | 83 | F | 28 | 4 | Yes | Yes |
| NP21-00057 | PD | SN | 83 | F | 28 | 4 | Yes | Yes |
| NP21-00057 | PD | AMY | 83 | F | 32 | 4 | Yes | Yes |
| NP21-00208 | PD | PFC | 81 | M | 87 | 6 | Yes | Yes |
| NP21-00208 | PD | PUT | 81 | M | 87 | 6 | Yes | Yes |
| NP21-00208 | PD | SN | 81 | M | 87 | 6 | Yes | Yes |

| | | | | | | | | |
|---|---|---|---|---|---|---|---|---|
| NP21-00208 | PD | AMY | 81 | M | 87 | 6 | Yes | Yes |
| NP21-00217 | PD | PFC | 84 | F | 43 | 6 | Yes | Yes |
| NP21-00217 | PD | PUT | 84 | F | 43 | 6 | Yes | Yes |
| NP21-00217 | PD | SN | 84 | F | 43 | 6 | Yes | Yes |
| NP21-00217 | PD | AMY | 84 | F | 32 | 6 | Yes | Yes |
| NP22-00037 | Control | AMY | 90 | M | 56 | 0 | Yes | Yes |
| NP22-00037 | Control | SN | 90 | M | 56 | 0 | Yes | Yes |
| NP22-00037 | Control | PUT | 90 | M | 56 | 0 | Yes | Yes |
| NP22-00037 | Control | PFC | 90 | M | 56 | 0 | Yes | Yes |
| NP22-00055 | PD | PFC | 76 | F | 59 | 6 | Yes | Yes |
| NP22-00055 | PD | PUT | 76 | F | 59 | 6 | Yes | Yes |
| NP22-00055 | PD | SN | 76 | F | 59 | 6 | Yes | Yes |
| NP22-00055 | PD | AMY | 76 | F | 59 | 6 | Yes | Yes |
| NP22-00075 | Control | AMY | 74 | M | 115 | 0 | No - hypoxic | Yes |
| NP22-00075 | Control | SN | 74 | M | 115 | 0 | No - hypoxic | Yes |
| NP22-00075 | Control | PUT | 74 | M | 115 | 0 | No - hypoxic | Yes |
| NP22-00075 | Control | PFC | 74 | M | 115 | 0 | No - hypoxic | Yes |
| NP23-00021 | PD | AMY | 71 | M | 114 | 5 | Yes | Yes |
| NP23-00021 | PD | SN | 71 | M | 114 | 5 | Yes | Yes |
| NP23-00021 | PD | PUT | 71 | M | 114 | 5 | Yes | Yes |
| NP23-00021 | PD | PFC | 71 | M | 32 | 5 | Yes | Yes |
| P73 | PD | SN | 92 | M | 32 | 5 | Yes | |
| P73 | PD | PFC | 92 | M | 32 | 5 | Yes | |
| P74 | PD | SN | 63 | M | 32 | 4 | Yes | |
| P74 | PD | PFC | 63 | M | 32 | 4 | Yes | |
| PT231 | Control | SN | 84 | M | 32 | 0 | Yes | |
| PT231 | Control | PFC | 84 | M | 32 | 0 | Yes | |

***Table S2. Mean variance explained by individual covariates.*** *Mean variance explained by individual covariates (design = ~ batch + sex + condition) using all features (HERVs, L1s, and genes) as calculated by fitExtractVarPartModel (variancePartition).*

| Pseudobulk (region_cluster) | Mean variance from sequencing batch | Mean variance from sex |
|---|---|---|
| AMY_0 | 0,39231496 | 0,1076047 |
| AMY_1 | 0,41895756 | 0,06247276 |
| AMY_2 | 0,21175977 | 0,04571372 |
| AMY_3 | 0,28960846 | 0,04988992 |
| AMY_4 | 0,32576012 | 0,09084974 |
| AMY_5 | 0,13742857 | 0,06148692 |
| PFC_0 | 0,31311396 | 0,05723861 |
| PFC_1 | 0,31918144 | 0,03531423 |
| PFC_2 | 0,23974501 | 0,02845113 |
| PFC_3 | 0,42109326 | 0,06623028 |
| PFC_4 | 0,53282174 | 0,15830616 |
| PFC_5 | 0,22311288 | 0,05055917 |
| PUT_2 | 0,1813245 | 0,0343387 |
| PUT_3 | 0,17805116 | 0,03907945 |
| PUT_4 | 0,25637325 | 0,05367004 |
| PUT_5 | 0,13092601 | 0,02670692 |
| SN_2 | 0,18091526 | 0,04250906 |
| SN_3 | 0,2640864 | 0,05187649 |
| SN_4 | 0,20861184 | 0,0481163 |
| SN_5 | 0,17807142 | 0,03279578 |
